# The *Acinetobacter baumannii* Mla system and glycerophospholipid transport to the outer membrane

**DOI:** 10.1101/384685

**Authors:** Cassandra Kamischke, Junping Fan, Julien Bergeron, Hemantha D. Kulasekara, Zachary D. Dalebroux, Anika Burrell, Justin M. Kollman, Samuel I. Miller

## Abstract

The outer membrane (OM) of Gram-negative bacteria serves as a selective permeability barrier that allows entry of essential nutrients while excluding toxic compounds, including antibiotics. The OM is asymmetric and contains an outer leaflet of lipopolysaccharides (LPS) or lipooligosaccharides (LOS) and an inner leaflet of glycerophospholipids (GPL). We screened *Acinetobacter baumannii* transposon mutants and identified a number of mutants with OM defects, including an ABC transporter system homologous to the Mla system in *E. coli*. We further show that this opportunistic, antibiotic-resistant pathogen uses this multicomponent protein complex and ATP hydrolysis at the inner membrane to promote GPL export to the OM. The broad conservation of the Mla system in Gram-negative bacteria suggests the system may play a conserved role in OM biogenesis. The importance of the Mla system to *Acinetobacter baumannii* OM integrity and antibiotic sensitivity suggests that its components may serve as new antimicrobial therapeutic targets.

## INTRODUCTION

Gram-negative bacteria are enveloped by two lipid bilayers, separated by an aqueous periplasmic space containing a peptidoglycan cell wall. The inner membrane (IM) is a symmetric bilayer of glycerophospholipids (GPL), of which zwitterionic phosphatidylethanalomine (PE), acidic phosphatidylglycerol (PG), and cardiolipin (CL) are among the most widely distributed in bacteria (1). In contrast, the outer membrane (OM) is largely asymmetric and composed of an inner leaflet of GPL and an outer leaflet of lipopolysaccharide (LPS) or lipooligosaccharide (LOS) (2). The OM forms the first line of defense against antimicrobials by functioning as a molecular permeability barrier. The asymmetric nature of its lipid bilayer and the structure of LPS/LOS molecules facilitates barrier function, as the core region of LPS impedes the entry of hydrophobic molecules into the cell while the acyl chains within the bilayer also help to prevent the entry of many hydrophilic compounds (3). Although progress has been made in understanding many aspects of OM assembly – including the discovery of an LPS transport system and the machinery for proper folding and insertion of outer membrane proteins (4, 5) – little is known about the molecular mechanisms for transport of the GPLs necessary for OM formation and barrier function.

*Acinetobacter baumannii* is an important cause of antibiotic-resistant opportunistic infections and has significant innate resistance to disinfectants and antibiotics. Similar to other Gram-negative opportunistic pathogens such as *Pseudomonas aeruginosa* and *Klebsiella* spp., individuals with breached skin or damaged respiratory tract mucosa are most vulnerable (6, 7). We performed a genetic screen to identify genes important for the OM barrier of *A. baumannii*. This led to the identification of an ABC (<u>A</u>TP-<u>b</u>inding <u>c</u>assette) transporter complex that promotes GPL export to the OM. Transporter disruption attenuates bacterial OM barrier function, resulting in increased susceptibility of *A. baumannii* to a wide variety of antibiotics.

The homologous system for *E. coli* has previously been termed Mla for its suggested role in the <u>m</u>aintenance of outer membrane <u>l</u>ipid <u>a</u>symmetry via the removal of GPL from the outer leaflet of the OM to the IM. While this is a reasonable hypothesis, there is not direct biochemical evidence that the Mla system functions to return GPL from the OM to the IM. In this work, we present evidence that the *A. baumannii* Mla system functions to promote GPL movement from the IM to the OM. This conclusion is based on the observation that newly synthesized GPLs accumulate at the IM of *mla* mutants, akin to how LPS molecules accumulate at the inner membrane in bacteria with mutations in the *lpt* genes encoding the LPS ABC transport system (5). Given the broad conservation of Mla in prokaryotic diderm organisms, the anterograde trafficking function of Mla might be exploited by a variety of species.

## RESULTS

### A screen for activity of a periplasmic phosphatase identifies genes required for *A. baumannii* OM barrier function

We identified strains with mutations in genes required for maintenance of the *Acinetobacter baumannii* OM barrier by screening transposon mutants for the development of a blue colony phenotype on agar plates containing the chromogenic substrate BCIP-Toluidine (XP). Although *A. baumannii* carries an endogenous periplasmic phosphatase enzyme, colonies remain white on agar plates containing XP. We reasoned that lesions in genes necessary for the OM barrier function should result in a blue colony phenotype, as the XP substrate becomes accessible to the periplasmic enzyme (8, 9). Screening roughly 80,000 transposon-containing colonies for the blue colony phenotype yielded 364 blue colonies whose insertions were mapped to 58 unique genes (Table S1). We confirmed the results of the screen by assaying for OM-barrier defects using ethidium bromide (EtBr) and N-Phenyl-1-naphthylamine (NPN) uptake assays (10, 11). We also tested for resistance to antimicrobials, including trimethoprim, rifampicin, imipenem, carbenicillin, amikacin, gentamicin, tetracycline, polymyxin B, and erythromycin. Greater than 85% of the strains identified in the screen demonstrated decreased OM barrier function compared to wild type. Out of the 58 strains with transposon insertions, 23 demonstrated OM permeability defects by NPN and EtBr uptake assays, and 49 out of 58 resulted in increased sensitivity to two or more antibiotics compared to the parent strain, indicating that the screen identified lesions causing OM barrier defects leading to increased permeability to small charged and hydrophobic molecules, including commonly used antibiotics.

### The Mla system is necessary for *A. baumannii* OM integrity

Four mutants with a blue colony phenotype contained unique transposon insertions in the genetic loci A1S_3103 and A1S_3102, predicted to encode core components (*mlaF* and *mlaE*) of a multicomponent ABC transport system. These genes are within a five-gene operon that encodes for a conserved proteobacterial ABC transport system homologous to the *E. coli mla* system previously implicated in OM integrity (12). The *A. baumannii* operon includes: *mlaF* and *mlaE*, respectively predicted to encode the nucleotide-binding and transmembrane domains of an ABC transporter; *mlaD*, encoding a protein containing an IM-spanning domain and a predicted periplasmic soluble domain; *mlaC*, encoding a soluble periplasmic protein; and *mlaB*, an additional gene predicted to encode a cytoplasmic <u>s</u>ulfate <u>t</u>ransporter and <u>a</u>nti-<u>s</u>igma factor antiagonist (STAS)-domain protein (Fig. 1A). An additional putative OM lipoprotein, is encoded on *mlaA*, or *vacJ*, which is clustered with the rest of the *mla* operon in some Gram-negative bacteria, although it is at a different chromosomal location in *A. baumannii*. MlaA has been functionally associated with the rest of the Mla components in *E. coli*, as mutations in *mlaA* yield comparable phenotypes to mutations in other components of the system (8).

**Fig. 1.**
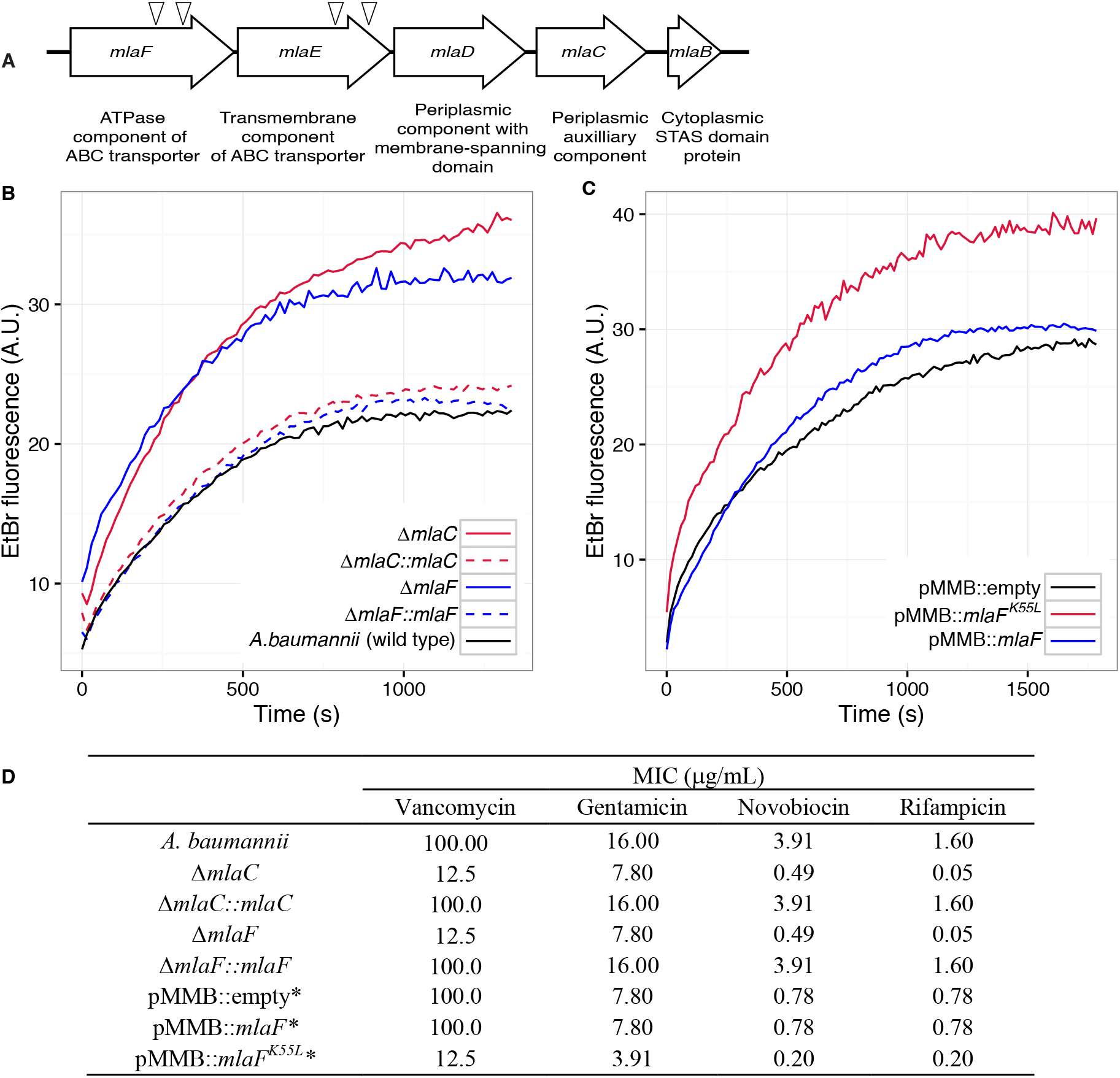
Disruption of the Mla system results in an altered outer membrane barrier. **(A)** Genomic organization of the *A. baumannii mlaFEDCB* operon and its predicted products. Triangles indicate the position of four independent transposon insertions, isolated in a screen for genes involved in outer membrane integrity. **(B)** Ethidium bromide uptake assay of outer membrane permeability of Δ*mla* mutants and complemented strains. A.U., arbitrary units. Lines shown depect the average of three technical replicates. **(C)** Ethidium bromide uptake assay of outer membrane permeability following plasmid-based expression of MlaF, compared to its dominant negative version, MlaF^K55L^. Lines shown depict the average of three technical replicates. **(D)** Minimum inhibitory concentration (MIC) of select antibiotics in *A. baumannii*. *Indicates wild type *A. baumannii* containing pMMB plasmid constructs, and cultures grown with the addition of kanamycin (25 μg/mL) to maintain plasmids and 50 μM IPTG for induction.

Bioinformatic analysis predicts that the *mlaC* and *mlaF* genes respectively encode the soluble periplasmic component and cytoplasmic ATPase component of the ABC transport system, and we chose to focus on mutants of these genes for further experiments to elucidate the function of the *mlaFEDCB* operon. Chromosomal deletions were created by allelic exchange, and these mutations resulted in OM permeability defects as measured by EtBr uptake assays. We complemented the OM defect for the Δ*mlaC* and Δ*mlaF* deletion mutants by repairing the original deletion event in the chromosome, and confirmed complementation of the observed permeability defect (Fig. 1B). Deletions in *mlaF* and *mlaC* also rendered *A. baumannii* increasingly sensitive to a variety of antibiotics as determined by MIC measurements (Fig. 1D). Increased sensitivity to antibiotics whose uptake is not mediated by OM porins is consistent with a direct effect on the membrane component of the OM permeability barrier (13, 14). In addition to OM defects, the *mla* mutants display phenotypes that may correlate with OM stress, including increased production of extracellular carbohydrates as evidenced by crystal violet staining of pellicles following growth in broth culture (Fig. S1A). These data indicate a role for Mla in the maintenance of the outer membrane barrier of *A. baumannii*.

### ATPase activity of MlaF is required for maintenance of the OM barrier of *A. baumannii*

To exclude the possibility that the membrane defect was the result of the disruptive effect of a partially formed Mla protein complex, we engineered an enzymatically inactive ATPase component and expressed the defective enzyme from a plasmid. We reasoned that by expressing this allele in the wild type bacteria we could create a dominant-negative effect on Mla function. The cytoplasmic ATPase component of the Mla system, MlaF, contains the consensus sequence GxxxxGKT at residues 49-56, characteristic of a Walker A motif. Downstream residues 173-178 contain the sequence LIMYDE, typical of a Walker B motif. The Walker motifs form highly conserved structures critical for nucleotide binding and hydrolysis (15). The lysine residue of the Walker A motif is particularly essential for the hydrolysis of ATP. Mutations in this lysine residue are inhibited for nucleotide binding, and the mutated protein is rendered inactive (16). Additionally, ATPase mutants in the key lysine residue have been shown to have a dominant-negative effect on ATP hydrolysis when co-expressed with their wild-type ATPase counterparts, as typical ABC transporters have a structural requirement for two functional nucleotide-binding proteins which dimerize upon substrate transport (17, 18).

Therefore, we created a version of the MlaF coding sequence with a leucine substitution of the Walker A lysine residue (MlaF^K55L^), and then cloned the mutated *mlaF* into the low-copy pMMBkan vector under control of the *mlaF* native promoter. We observed that expression of MlaF^K55L^ in wild type *A. baumannii* had a dominant-negative effect on membrane permeability as measured by EtBr uptake (Fig. 1C), and expression of MlaF^K55L^ also resulted in increased exopolysaccharide production as demonstrated by increased staining by crystal violet (Fig S1B). Correspondingly, expression of MlaF^K55L^ rendered *A. baumannii* more sensitive to a variety of antibiotics, resulting in reduced MICs when compared to *A. baumannii* expressing the empty pMMBkan vector (Fig. 1D). Therefore, expression of a defective ATPase results in a dominant-negative mutant with a comparable phenotype to deletion of components of the *mla* operon.

These results demonstrate a requirement for ATP hydrolysis by MlaF for the maintenance of OM barrier function in *A. baumannii*, and indicate that the phenotypes of deletion mutants were likely a result of a lack of transport function, rather than formation of a toxic incomplete membrane protein complex.

### Structure of the *A. baumannii* MlaBDEF complex

The genetic arrangement and conservation of the components of this ATPase-containing transport complex indicated it was likely that the individual components formed a higher order protein structure. To define whether the Mla components form a stable protein complex, we expressed the entire operon (*mlaFEDCB*) from *A. baumannii* ATCC 17978 in *E. coli* with a carboxy-terminal hexahistidine tag on the MlaB component. Affinity purification of MlaB revealed three additional bands, with sizes corresponding to MlaF, MlaD, and MlaE (Fig. S2) and confirmed by MALDI-TOF mass spectrometry analysis, indicating that these four proteins form a stable complex. We did not detect MlaC, suggesting it might interact only transiently with the other components, consistent with results recently reported by Thong et al. (19).

We next used cryo-electron microscopy to characterize the architecture of the *A. baumannii* MlaBDEF complex (abMlaBDEF). This complex is uniformly dispersed in vitreous ice (Fig. S3A), and 2D classification demonstrated the presence of a range of views suitable for structure determination (Fig. S3B). Following 2D- and 3D-classification, we obtained a final dataset of ~ 14,000 particles with which we obtained a structure to a resolution of 8.7 Å (Fig. S3D). The structure possesses significant visible features in agreement with the nominal resolution (Fig. S3C). Based on the bioinformatically-predicted localization of individual proteins and work recently performed on the similar *E. coli* Mla complex (ecMlaBDEF) (19), we propose that MlaD localizes to the periplasmic side of the IM, MlaE forms the central transmembrane region, and MlaF and MlaB form the bottom layer on the cytoplasmic face of the IM with two visible hetero-dimers (Fig. S3E). We note that the structure of ecMlaBDEF, at lower resolution, was reported recently (20). The overall features of both structures, solved independently, are identical, suggesting that they correspond to the correct structure for the complex. However, the limited resolution of the ecMlaBDEF complex structure did not allow modeling of its individual subunits, in contrast to the abMlaBDEF structure reported here.

We note that a clear six-fold symmetry is present for the region of the map attributed to MlaD (Fig. 2B), despite the fact that we only imposed a 2-fold symmetry averaging. This agrees with the proposed hexameric state of its *E. coli* homologue (ecMlaD) (19). We next modeled abMlaD, using an evolution restraints-derived structural model of ecMlaD (21) as a template, and used our previously-published EM-guided symmetry modeling procedure (22) to model its hexameric state. The obtained abMlaD hexameric model is at a low-energy minimum (Fig. S4B) and fits the EM map density well (Fig 2B and S5B Fig). A crystal structure of the periplasmic domain of ecMlaD published recently (20) formed a crystallographic hexamer, suggesting that this corresponds to the native hexomeric arrangement for this domain. Our abMlaD hexameric model is very similar to the crystallographic ecMlaD structure (Fig. S4C), supporting the proposed domain arrangement in the MlaBDEF complex. We note, however, that one region of density in the EM map is not accounted for by our MlaD hexamer model (Fig. 2B). The localization of this extra density suggests that it corresponds to a ~ 45 amino-acid insert present between strands 4 and 5 of the abMlaD β-sheet (Fig. S5A). The role of this insert, uniquely found in the *A. baumannii* orthologue, is not known.

**Fig. 2.**
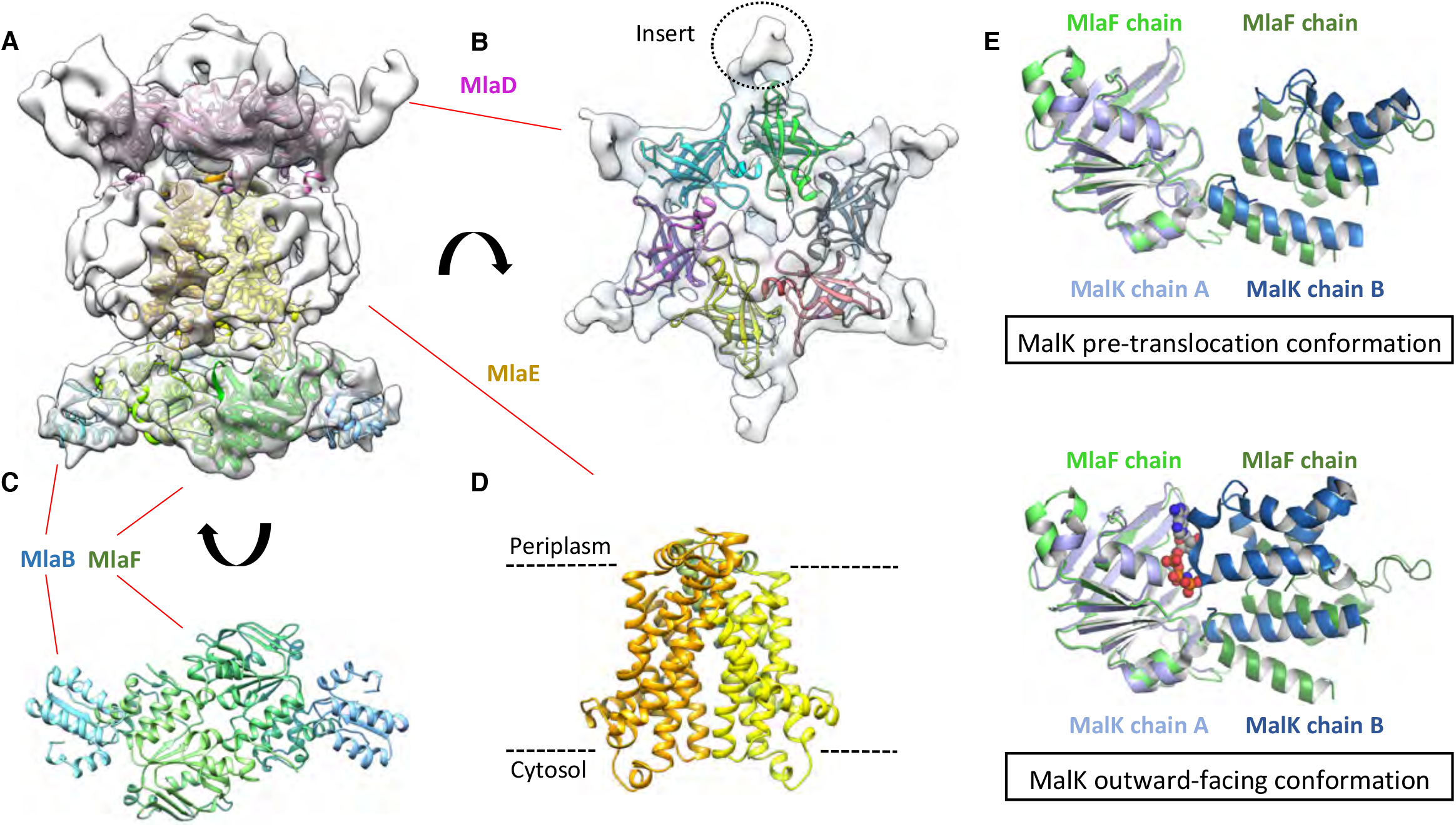
Structure of the abMlaBDEF complex. (A) Cryo-EM mal of abMlaBDEF (grey), with structural models for MlaD, MlaE, MlaB and MlaF (in magenta, yellow, cyan and green respectively) docked at their putative location, as viewed from the side and bottom. The density for most helices is clearly resolved. (B) Cartoon representation of the MlaD hexameric model, with each subunit in grey, orange, yellow, magenta, cyan and green respectively. The putative position of the abMlaD insert is shown. (C) Cartoon representation of the MlaB and MlaF dimeric model, with MlaB in light and dark cyan and MlaF in light and dark green. (D) Cartoon representation of the MlaE dimeric model (in orange and yellow). (E) Comparison of the MlaF domain arrangement in the EM map to that of the Maltose transporter ATPase MalK. The two chains of MlaD (in light and dark green) superimpose well to those of MalK (in cyan and dark blue) in the pre-translocation conformation (left, PDB ID: 4KHZ), while a clear rotation is observed compared to the ATP-bound outward-facing conformation (right, PDB ID: 4KI0).

We next modeled the structures of MlaB and MlaF and fitted their respective coordinates in the corresponding region of the EM map (Fig. 2C and Fig. S4A). For both proteins, most helices are well resolved, which allowed us to place the models unambiguously. We then compared the conformation of the ATPase MlaF to that of the maltose transporter ATPase MalK, which has been trapped in several conformations of the transporter; i.e. the inward-facing state, the pre-translocation state, and the outward-facing state (23, 24). Interestingly, the arrangement of MlaF clearly resembles the pre-translocation state of MalK (Fig. 2D). This suggests that we have trapped a similar conformation of the abMlaBDEF complex. It is possible that MlaD and/or MlaF, for which there are no equivalent in other ABC transporters, stabilizes this conformation. Alternatively, it is possible that the presence of detergents, which were present to solubilize the complex, mimics the natural ligand in the transporter’s active site. Finally, the transmembrane (TM) region of the map is well resolved, and density for the transmembrane (TM) helices can be clearly identified. We therefore modeled abMlaE, using an evolution restraints-derived structural model of ecMlaE (21) as a template, and fitted the obtained coordinates in the corresponding region of the map, with the orientation corresponding to the predicted topology. The resulting MlaE dimer model (Fig. 2D) fits well to the EM map density (Fig. S5C), and clearly corresponds to a closed transporter, with no solvent channel between the subunits. Interestingly, we also noted clear density for three TM helices that likely correspond to the MlaD N-terminal helices (Fig. 3A). However, they lacked continuity, and we observed that only two form a direct interaction with MlaE. It is possible that this is due to heterogeneity in the orientation of MlaD relative to the rest of the complex. To verify this, we performed further 2D classification of the particles used for reconstruction (Fig. 3B), which revealed a range of positions for the MlaD region relative to the rest of the complex. We therefore performed further 3D classification leading to a smaller dataset of ~ 8,000 particles. This produced a structure of lower resolution (~ 11.5 Å) but with the six MlaD N-terminal TM helices clearly visible (Fig. 3B). While the periplasmic domain possesses 6-fold symmetry, the TM domains of MlaD do not appear symmetrical, with two forming close contacts with the density attributed to MlaE while the other four do not appear to contact any other proteins. This observation likely explains the asymmetry of contacts between the dimeric MlaE and the hexameric MlaD. A higher-resolution structure will be required to determine if additional contacts are formed between the outward-facing loops of MlaE and the periplasmic domain of MlaD.

**Fig. 3.**
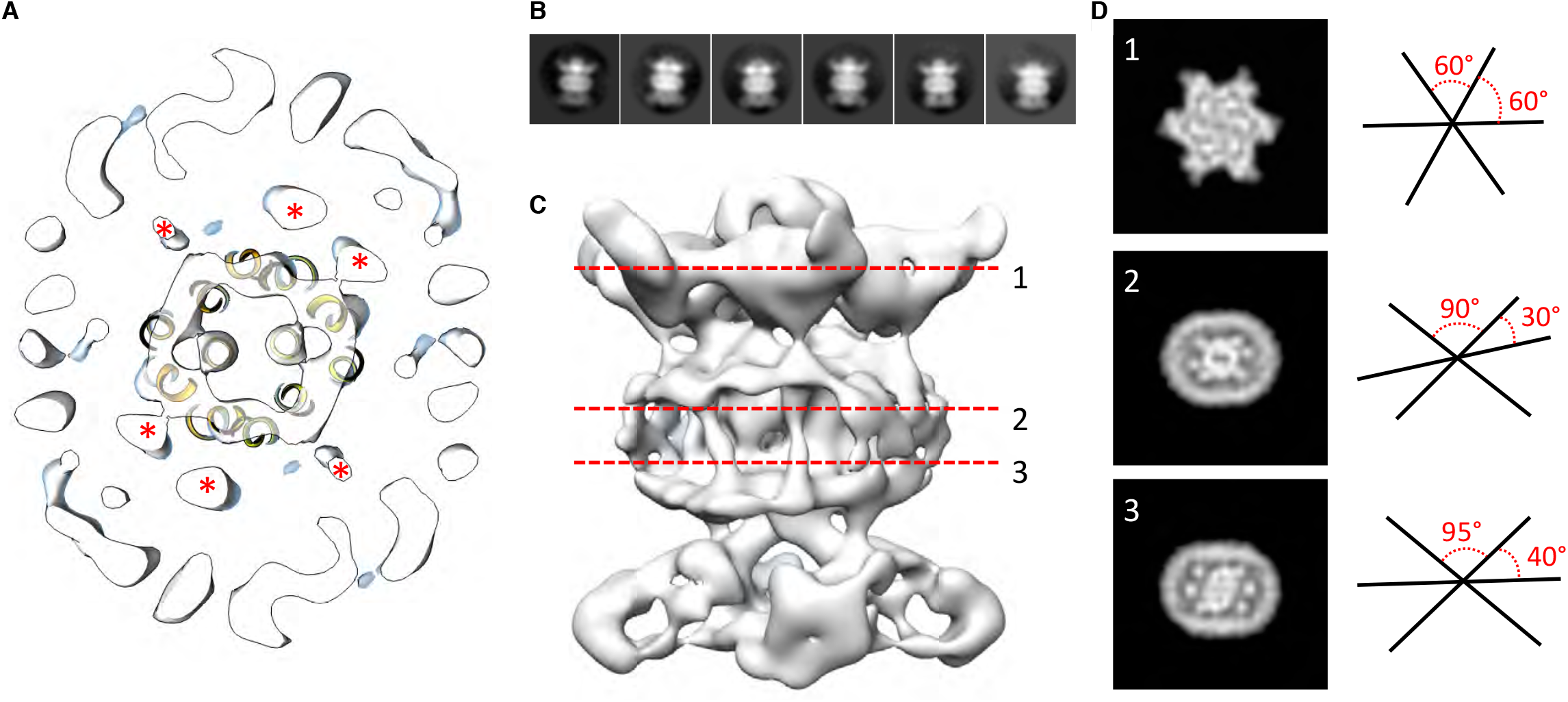
Localization of the 6 TM helices from MlaD. **(A)** lateral section of the abMlaBDEF EM map, with the MlaE model in yellow. Density attributed to the MlaD N-terminal helices are indicated with a red star. **(B)** Reference-free 2D classes generated from the set of particles used to generate the abMlaBDEF structure, corresponding to side views. A range of orientations for the periplasmic domain is observed. **(C)** Structure of abMlaBDEF, generated using a subset of the most homogenious ~8,000 particles. Somefeatures of the map shown in 3 A are not present, but the overall structure is similar. Six well-defined helices in the central TM region are visible. **(D)** Sections along the vertical axis, corresponding to the three red lines shown in B, is shown on the left. The six-fold axis of MlaD is visible in the periplasmic region, but this breaks down in the TM region, where the six helices are asymmetric. An angular representation of the six helices is represented on the right.

### Components of the Mla system interact directly with GPL

The crystal structure of MlaC has been solved from *Ralstonia solanacearum*. The structure contains a single phosphatidylethanolamine molecule oriented such that the hydrophobic acyl chains are located inside the protein while the hydrophilic head group is exposed (25). More recently, the crystal structure for MlaC has been solved from *E. coli* and shown to bind lipid tails (20). As noted in previous work performed on the *E. coli* Mla system, this is strong evidence that the substrates of the Mla system are GPL (12). In order to confirm that the periplasmic components of the Mla pathway in *A.baumannii* interact with GPL, we purified the soluble domains of both MlaC and MlaD by expressing histidine-tagged proteins followed by Ni-affinity FPLC purification. After overnight dialysis of the proteins, we performed Bligh-dyer chloroform extraction on the purified proteins to isolate any bound GPL and analyzed the results by LC-MS/MS. GPL analysis revealed both phosphatidylglycerol and phosphatidylethanolamine of varying acyl chain lengths. This suggests the possibility that the periplasmic substrate binding components of the system may bind diacylated GPL molecules with limited polar head group specificity (Fig. S6).

### Mla mutants have decreased abundance of outer membrane GPL

Given the OM defect of *mla* mutants, as well as the system’s apparent direct association with GPL, we chose to further characterize the overall membrane GPL composition of the *mla* mutants. Previous work on the Mla system in *E.coli* has demonstrated an increase in hepta-acylated lipid A in *mla* mutants, indicating activation of PagP that acylates GPL and lipid A in the outer leaflet of the OM in enterobacteria (12, 26). From this data it has been suggested that the system may serve to maintain lipid asymmetry within the OM, although it is well known that GPL displacement to the OM outer leaflet is a general reflection of chemical damage to the OM (27-29). However, biochemical analysis of the membrane GPL composition for *mla* mutants has not been published for any organism to our knowledge, so we sought to apply our lab’s methods of GPL quantification to test the hypothesis of retrograde transport function. To determine whether *A. baumannii mla* mutations cause changes in the membrane GPL concentration, GPL were extracted from inner and outer membrane fractions separated by density centrifugation. Thin-layer chromatography (TLC) and electrospray-ionization time-of-flight mass spectrometry (ESI-MS) were used to qualitatively assess GPL composition. TLC and ESI-MS indicated Δ*mlaC A. baumannii* had a dramatically decreased abundance of all major phospholipid species in the OM compared to wild type. (Fig. 4A and Fig. S7).

**Fig. 4.**
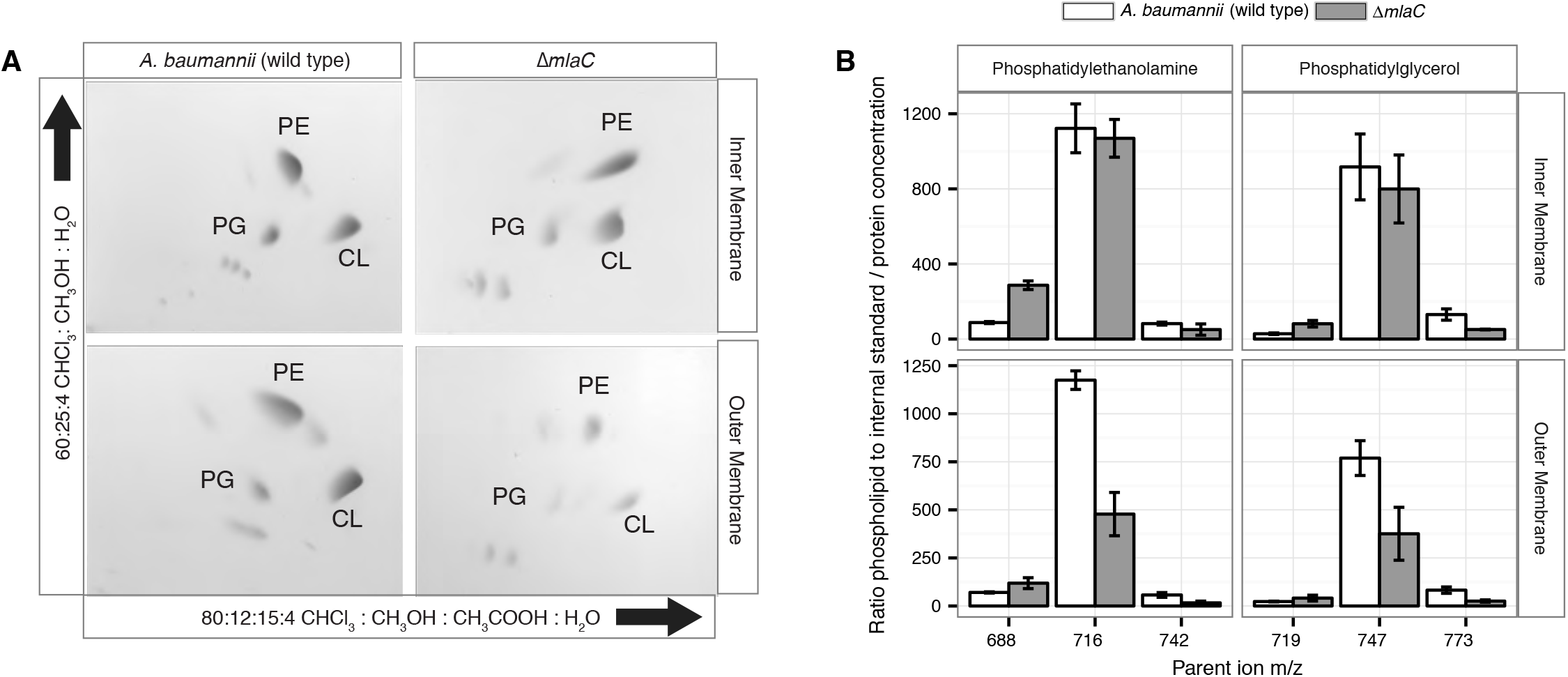
Outer membrane glycerophospholipid levels are reduced in Δ*mlaC* mutant. **(A)** Identification of inner and outer membrane phospholipids of wild type *A. baumannii* and Δ*mlaC* using 2D thin-layer chromatography. PE, phosphatidylethanolamine; PG, phosphatidylglycerol; CL, cardiolipin. **(B)** LC-MS/MS quantification of isolated inner and outer membrane glycerophospholipids. Error bars indicate ± s.e.m. (n = 3).

To better analyze the differences in membrane GPL, we quantified GPL by normal phase liquid-chromatography collision-induced-dissociation mass spectrometry (LC-MS/MS). We quantified the ratio of individual GPL within each membrane by normalizing to an internal standard of known quantity. We then normalized the quantified GPL to the protein content of isolated IM and OM. Quantitative LC-MS/MS confirmed the overall reduction in outer membrane GPLs observed by ESI-MS and TLC, with the reduced levels observable across multiple GPL species for Δ*mlaC* mutants relative to wild type (Fig. 4B). Therefore, mutations in the components of the Mla system result in a decrease in OM GPL, whereas the retrograde transport hypothesis would predict an increase in OM GPL. Therefore, these results instead suggest a possible role for Mla in outward GPL trafficking.

### Mla mutants demonstrate an accumulation of newly synthesized GPL in the IM

The overall decrease in outer membrane glycerophospholipids of A. baumannii *mla* mutants suggests that either the Mla system is functioning to deliver GPLs from the inner membrane to the outer membrane, or alternatively, mutations in the Mla system may disrupt the outer membrane in a manner that leads to the activation of outer membrane phospholipases, which then degrade GPL. Work performed on the Mla system in *E.coli* has demonstrated that disruption of genes in the Mla pathway results in activation of both the OM acyl-transferase PagP, which cleaves a palmitate moiety from GPL and transfers it to LPS and PG, creating a hepta-acylated LPS molecule and palmitoyl-PG and the OM phospholipase PldA (12, 28). *A. baumannii* has no known PagP enzyme but similar activity of the multiple predicted OM phospholipases could account for the reduction in OM GPL as observed by TLC and quantitative mass spectrometry. Therefore, we designed a mass spectrometry-based assay to study intermembrane GPL transport using ^13^C stable isotope labeling (Fig. S8A), to better analyze the directionality of GPL transport by the Mla system between the bacterial membranes,. When grown in culture with sodium acetate as the sole carbon source, many bacteria directly synthesize acetyl-CoA using the conserved enzyme acetyl-CoA synthase (30). Acetyl CoA, the precursor metabolite for fatty acid biosynthesis, is first converted to malonyl-CoA and enters the FasII (fatty acid biosynthesis) pathway that supplies endogenously synthesized fatty acids to macromolecules such as lipopolysaccharides, phospholipids, lipoproteins, and lipid-containing metabolites. By growing cultures in unlabeled acetate then “pulsing” with 2-^13^C acetate and analyzing separated membrane fractions from set time points, we can observe the flow of newly synthesized GPLs between the IM and OM of *A. baumannii* (Fig. S8B) (26).

Upon introducing the 2-^13^C acetate as the sole carbon source, ^13^C-labeled GPL were immediately synthesized in the bacterial cytoplasm. We reasoned that continued growth in ^13^C acetate should result in a mixed pool of unlabeled and labeled IM GPL molecules. As the GPL are then fluxed from the IM to the OM, the likelihood that an individual GPL molecule is transported is directly proportional to the ratio of labeled to unlabeled GPL in the IM pool. As the bacteria continue to grow in ^13^C acetate, the ratio of labeled to unlabeled GPL in the IM will gradually increase as new GPL are synthesized and inserted in the IM. As such, the likelihood of transporting labeled GPL to the OM will also increase. A comparison of the ratios of labeled to unlabeled GPL in the IM and OM will thus reflect the efficiency of transport between the membranes, and analysis of transport in wild type *A. baumannii* will establish reference for transport efficiency with which to compare our mutants. Additionally, OM phospholipases, some of which may be activated upon membrane damage (31), will not distinguish between labeled and unlabeled GPL and therefore will not affect the ratio of labeled to unlabeled GPL obtained from this assay.

Membrane separation and analysis of wild type *A. baumannii* revealed near-identical rates-of-change between the two membranes in ratios of ^13^C-labeled to unlabeled GPLs, indicating that newly synthesized GPLs are transported and inserted into the OM at a rate equivalent to their rate of synthesis and assembly within the IM. Furthermore, the ratios of labeled to unlabeled GPLs were nearly equal in the IM compared to the OM at the time points evaluated indicating that GPL transport likely occurs rapidly, consistent with earlier pulse-chase experiments performed in *E. coli* that estimate the half-life of translocation of various GPLs at between 0.5 and 2.8 min (32). By contrast, mutants in the Mla system accumulate newly synthesized GPLs in their IM at a greater rate than occurs in the OM as evidenced by the increasing disparity in the ratio of labeled to unlabeled GPLs between the IM and OM over time (Fig 5A). The discrepancy in ratios of labeled to unlabeled GPLs between the IM and OM of Δ*mlaF* is apparent for PG and PE of varying acyl chain lengths corresponding to the most naturally abundant species C16:0/C16:0, C18:1/C18:1, or C16:0/C18:1 (Table S2). Further, the effects of MlaF^K55L^ expression on GPL trafficking were similar to what was observed in the Δ*mlaF* strain (Fig. 5B). Therefore, ATP hydrolysis by MlaF appears to be a requirement for extraction of these GPLs from the IM of *A. baumannii* for subsequent transport to the OM.

**Fig. 5.**
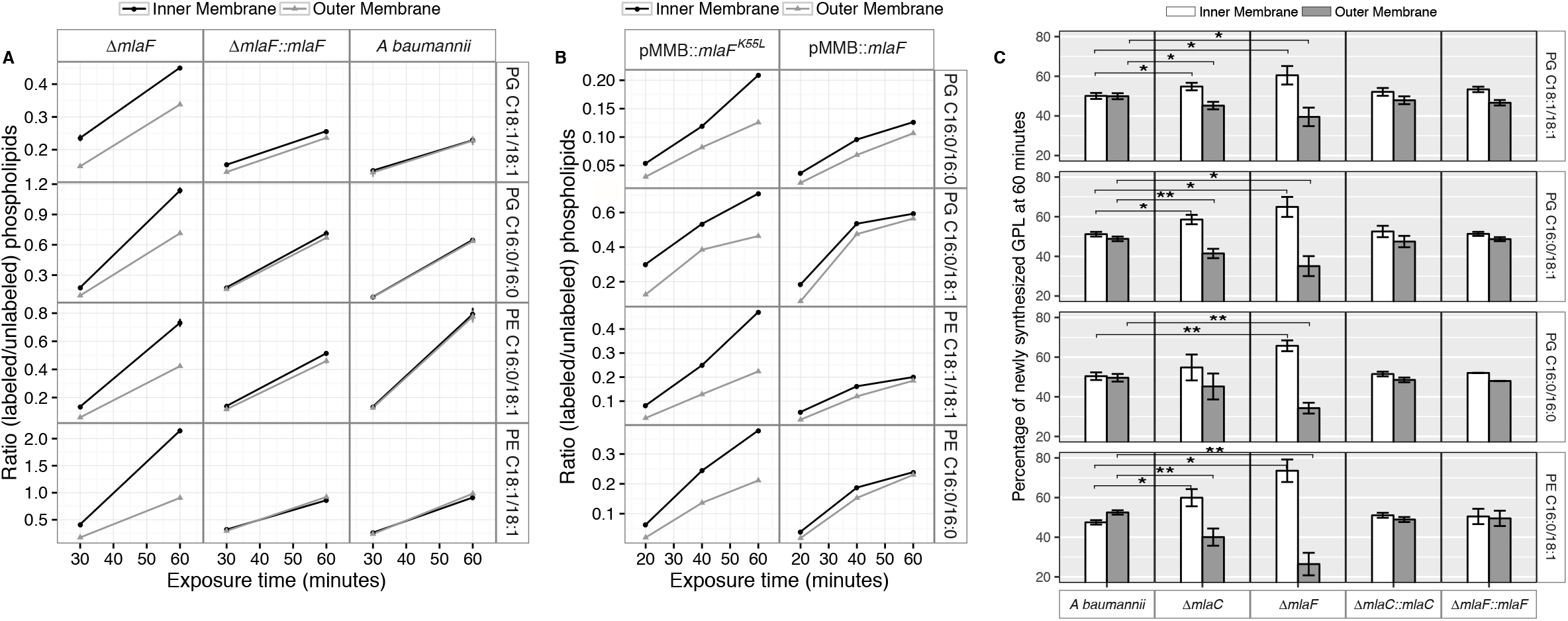
Newly synthesized glycerophospholipids accumulate at the inner membrane of Mla mutants. **(A)** LC-MS/MS quantification of ^13^C labelled/unlabeled glycerophospholipids in isolated membrane fractions over time after growth in 2-^13^C acetate in Δ*mlaF* and complemented strain. Facet labels on the right indicate the specific glycerophospholipid species analyzed and the acyl chain length. PG, phosphatidylglycerol; PE, phosphatidylethanolamine. Shown is representative data from repeated experiments. **(B)** LC-MS/MS quantification of ^13^C labelled/unlabeled glycerophospholipids in isolated membrane fractions following plasmid-based expression of MlaF compared to its dominant negative version, MlaF^K55L^. Facet labels on the right indicate the specific glycerophospholipid species analyzed and the acyl chain length. PG, phosphatidylglycerol; PE, phosphatidylethanolamine. Shown is representative data from repeated experiments. **(C)** Relative proportion of newly synthesized GPL on IM and OM after one hour growth in 2-^13^C acetate. Error bars represent ± s.d. (n = 2). Statistical analyses performed using a Student’s *t* test.p-Value: *, *p* < 0.05; **, *p* < 0.01.

To better characterize the role of the periplasmic substrate binding component MlaC, we performed similar stable isotope pulse experiments to observe the flow of newly synthesized GPLs in the Δ*mlaC* strains. Stable isotope experiments on Δ*mlaC* mutants reveal IM accumulation of newly synthesized GPLs similar to the result in Δ*mlaF* mutants (Fig. S9A), indicating that in the absence of the periplasmic component GPLs are not efficiently removed from the IM by the remainder of the Mla system. We also sought to characterize the potential role of the putative OM-lipoprotein MlaA, which has been implicated as a component of the Mla system in *E. coli*. A chromosomal deletion strain of *mlaA* was created by allelic exchange, and complemented by expression of MlaA from a pMMB67EH-Kan plasmid. The results of the stable isotope pulse experiments in the Δ*mlaA* strain revealed results consistent with those obtained from Δ*mlaC* and Δ*mlaF*, in which the ratio of labeled to unlabeled GPL is consistently higher in the inner membrane than the outer membrane after one hour of exposure to ^13^C-acetate (Figure S9B and S9C). These results are consistent with a model in which the IM-localized ABC transporter complex MlaBDEF first transfers GPLs to the periplasmic binding protein MlaC, which then transports GPL to the OM, whereupon MlaA facilitates GPL insertion into the OM.

## DISCUSSION

We performed a screen to identify *A. baumannii* proteins that are essential for its OM barrier that led to the identification of an ABC transport system whose ATPase activity maintains OM barrier function. IM and periplasmic components of this system can be purified, bind GPLs, and assemble into a defined protein complex with significant symmetry, indicating that this system could function to transport GPLs from the IM to the OM. Consistent with the possibility that Mla functions as an anterograde transporter, the OM of mutants show an overall reduction of GPL along with an excess accumulation of newly synthesized GPL on the IM. Therefore, these results lead us to propose that the function of the *A. baumannii* Mla system is the trafficking of GPL from the IM, across the periplasm, for delivery to the outer membrane (Fig. 6). According to this model, ATP hydrolysis by MlaF provides the initial energy to extract GPL from the IM, while the substrate binding components MlaD and MlaC take up lipids for delivery to the OM. It has been observed by van Meer and colleagues that complete extraction of GPLs from the membrane bilayer requires a Gibbs free energy of ~100 kJ/mol (33, 34), whereas ATP contains just 30 kJ/mol of energy. To account for the energy difference, a hydrophobic acceptor molecule is proposed to allow the lipids to fully dissociate from the rest of the ABC transporter and facilitate complete removal from the bilayer. The GPL-binding component, MlaD, contains an IM spanning domain and is shown here, and in orthologous systems, to be in complex with the MlaE and MlaF proteins within the IM (20, 35). The close association of MlaD with the outer leaflet of the IM may allow it to extract lipids from the IM by hydrophobic interaction with the acyl chains after ATP hydrolysis by MlaF. Subsequent trafficking across the periplasm then involves the periplasmic GPL binding protein MlaC, which likely accepts GPL from MlaD and then carries them to the OM. We note however the observed effect of *mlaC* deletion on GPL accumulation in the IM, while statistically significant for most of the analyzed diacyl-glycerophospholipids, appears to be less than that of deletion of the ATPase component (Fig. 5C), suggesting that while MlaC may participate in transfer to the OM, there may be redundant mechanisms by which the IM complex can transport or remove IM GPL in the absence of MlaC. While the precise mechanism of GPL insertion into the OM is not yet known, work performed on the *E.coli* Mla system has shown that MlaC interacts with both the IM MlaFEDB complex, as well as with the putative OM lipoprotein MlaA, and our results support a role for MlaA in the function of the overall Mla system and delivery of GPL to the OM.

**Fig. 6.**
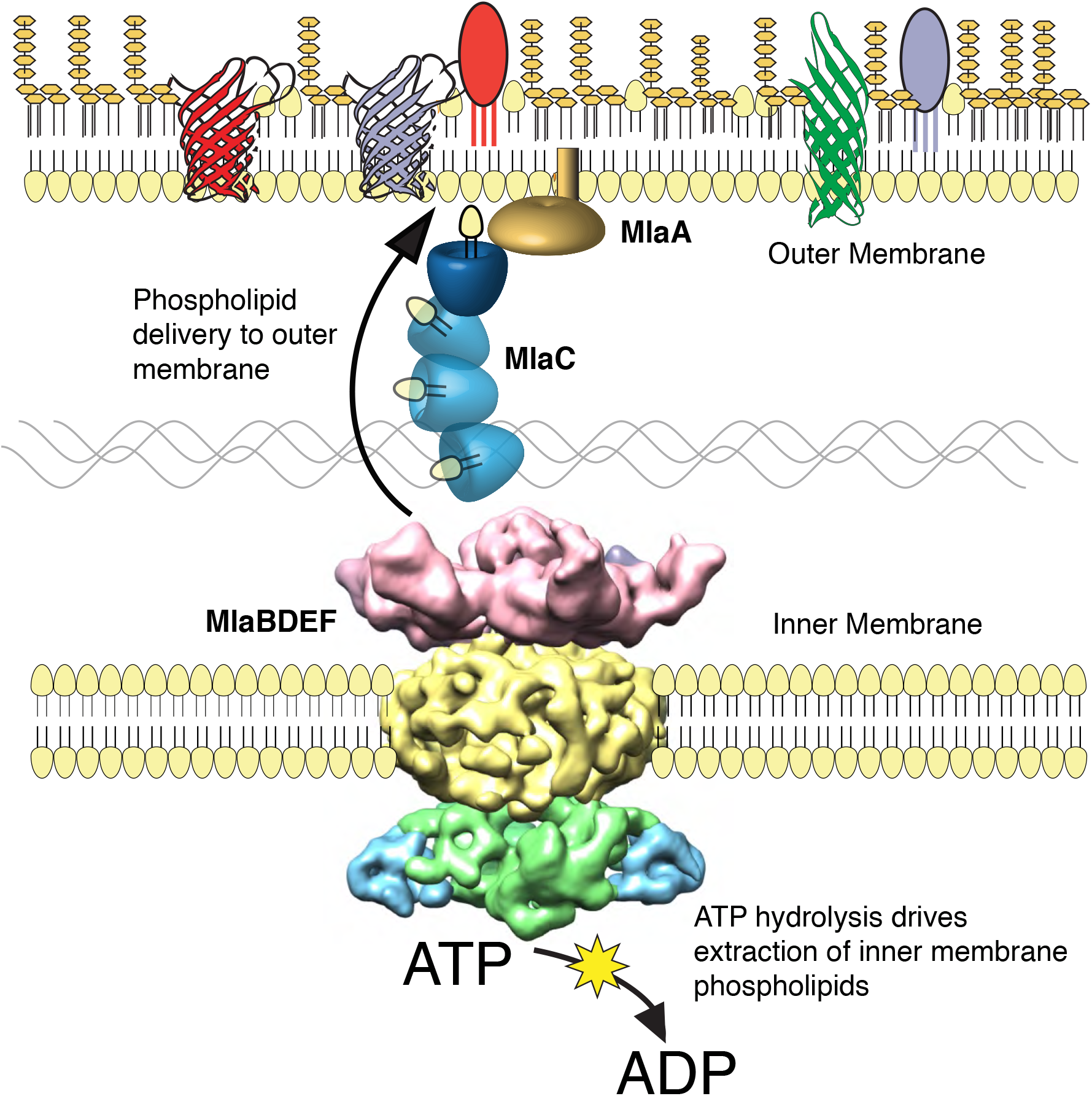
The multicomponent Mla system transports glycerophospholipids from the inner membrane to the outer membrane of *A. baumannii*. A schematic of glycerophospholipid transport to the Gram-negative bacterial outer membrane by the Mla system.

In this work, we designed a method to monitor lipid transport between Gram-negative bacterial membranes using stable ^13^C isotope labeling. Our results using this assay are consistent with the Mla system functioning as an anterograde GPL transporter, however they do not exclude the possibility of a dual role for Mla components in the maintenance of OM lipid asymmetry. Previous work performed on the orthologous Mla system in *E.coli* has been interpreted to suggest that the function of the system is to remove GPL from the outer leaflet of the OM for retrograde transport back into the cytoplasm based on the observation that *E.coli mla* mutants likely contain GPLs on the outer leaflet of the OM. (12, 36). This proposed function was inferred from the observation that gene deletions resulted in an increased activation of the OM-phospholipase enzymes PagP and OMPLA, suggesting an increased amount of GPL in the outer leaflet of the OM (12). The interpretation of retrograde transport function was also based on the existence of an orthologous system in plant chloroplasts that transports lipids from the endoplasmic reticulum (ER) into the organelle. Many plants require this retrograde transport function because certain lipids in the chloroplast thylakoid membrane derive from GPL originating in the ER (37). However, since Gram-negative bacteria synthesize GPL within the IM, retrograde transport of GPL would only be necessary for the recycling of GPL mislocalized to the OM outer leaflet. Although this is a reasonable inference based on data available at the time, we would point out that the directionality of transport by the *E. coli* Mla system had not been thoroughly probed experimentally using membrane analysis or with a functional assay of the type performed here. It is conceivable that the import function of the orthologous chloroplast TGD system is a result of adaptation to the intracellular environment, the system in this case having evolved to aid in the transfer of GPL from the nearby ER to the chloroplast. Furthermore, while it is possible that the Mla system in *E. coli* serves a different primary function than in *A. baumannii*, we demonstrate here that both complexes possess a similar architecture, pointing to a conserved function. The outer membrane defect phenotypes observed in *E. coli mla* mutants might also be explained by a disruption of OM structure stemming from decreased concentrations of OM GPL, leading to activation of the PagP enzyme. It is well established that for *E. coli*, GPL displacement to the OM outer leaflet and subsequent activation of these enzymes reflects OM instability and can be achieved by chemical disruption of the bilayer (27-29). It may be the case that the OM of *E. coli mla* mutants contain GPL in the outer leaflet, but the possibility remains that OM GPL can flip into the outer leaflet under conditions of OM damage resulting from an imbalance of LPS-to-GPL ratios, along with perhaps the corresponding disruption of OM proteins. However, final determination of the directionality of GPL transport by the Mla system in *E.coli* and other organisms will require intermembrane transport studies similar to what has been done here for *A. baumannii*, along with studies similar to those performed for the Lpt LPS transport system for which molecular transfer of LPS from molecule to molecule of the Lpt system is functionally defined.

The gene for MlaA, the proposed OM component, is at a different chromosomal location from the remainder of the *mla* operon. Recent structural studies on MlaA have revealed that MlaA forms a ring-shaped structure localized the inner leaflet of the OM, and have shown it to form stable complexes with the outer membrane proteins OmpF and OmpC (38). The proposed structure of MlaA in the OM supports the argument that MlaA is involved in removal of GPL from the outer leaflet, and it is suggested that GPL from the outer leaflet travel through a pore in MlaA where they are received by MlaC, yet our data reveals that *A. baumannii ΔmlaA* mutants are defective in delivery of GPL from the IM to the OM. These data can be reconciled by a model in which MlaA functions both to remove mislocalized GPL from the outer leaflet of the OM, and additionally serves to facilitate delivery of GPL to the OM by MlaC, perhaps by enabling MlaC localization to the surface of the inner leaflet. By this model, mutations in MlaA will be phenotypically similar to mutations in other components of the Mla system, and we would expect to observe a decreased rate of anterograde GPL transport. We would here point out that while previous work has implicated the Mla system in the maintenance of OM lipid asymmetry through observation of increased activity of PagP, the role of the MlaFEDB complex and MlaC in retrograde GPL transport has previously only been inferred from homology to the chloroplast TGD system. It is established that cellular mechanisms exist in Gram-negative bacteria to resist stressful conditions that lead to OM disruption. For example, OM phospholipase enzymes, such as PldA, are activated under conditions of membrane stress to digest GPL in the outer leaflet of the OM, as high levels of GPL in the outer leaflet destabilize the OM barrier function. The model of retrograde GPL transport by the Mla system proposes that growing cells expend cellular energy in the form of ATP in order to transport undigested GPL from the OM, across the periplasm, and back into the IM, at which point some of those same molecules will be transported back to the OM by an unknown mechanism. However, the available data points most clearly to a model of anterograde GPL transport by MlaFEDB and MlaC, facilitated in some way by MlaA.

The first three genes of the *mla* operon – comprising an ATPase, permease, and substrate-binding components of the ABC transporter complex – are conserved in *Mycobacteria spp, Actinobacteria*, and chloroplasts, while the entire five-gene operon appears to be conserved in Gram-negative bacteria (39). Given the conservation of the system across Gram-negative species, our results may shed light on a generalized mechanism contributing to OM biogenesis. Additionally, we have here demonstrated that the function of this ABC transport system is crucial for maintaining the integrity of the *A. baumannii* OM. The fact that *mla* mutations are tolerated, and that levels of OM GPL are reduced but not abolished, suggests the intriguing possibility of additional undiscovered mechanisms of GPL delivery to the OM. Also of interest is the potential role of the increased exopolysaccharide observed upon disruption of the Mla system. It is possible this exopolysaccharide plays a partially compensatory role in *A. baumannii* resulting from decreased OM GPL, given that recent work has shown that *A. baumannii* exopolysaccharides can contribute to antibiotic resistance, likely through improved barrier function (40).

The progression towards a more complete understanding of intermembrane GPL transport and OM barrier function should ultimately have relevance in the development of novel drug targets to undermine emerging antibiotic resistance in Gram-negative pathogens. The emergence of antibiotic resistant Gram-negative bacteria for which few or no antibiotics are available therapeutically is an important medical concern. This issue is typified by current isolates of *A. baumannii* that can only be treated with relatively toxic colistin antibiotics. This has led many individuals and agencies to propose the development of single agent antimicrobials which could be used for organisms such as *A. baumannii* and *P. aeruginosa* that have significant antibiotic resistance. Therefore, work furthering the understanding of the OM barrier could lead to the development of drugs which target the barrier and allow the therapeutic use of many current antibiotics.

## MATERIALS AND METHODS

### Bacterial strains

Transposon mutagenesis and subsequent chromosomal deletions of *mla* genes were performed in *Acinetobacter baumannii* ATCC 17978.

### A Mariner-based transposon vector for use in *Acinetobacter baumannii*

To perform transposon mutagenesis a Mariner-based transposon vector was designed for use in *Acinetobacter baumannii* ATCC 17978. The new transposon vector, derived from pBT20, termed pMarKT, contains an outward facing pTac promotor as well as a selectable kanamycin resistance marker followed by an omega terminator within the Mariner arm sites **(41)**. The plasmid backbone contains the Mariner transposase gene C9 Himar, a *tetRA* resistance marker from Tn10, a p15A origin from pACYC184, and an oriT site for mobilization. The plasmid was constructed by PCR of select fragments followed by restriction digest and ligation of the cleaved ends. The new transposon vector was confirmed by restriction digest and partial sequencing.

### Transposon mutagenesis

Initial mutagenesis revealed that many hits occurred in the high affinity phosphate uptake transcriptional repressor *phoU* (A1S_0256). Subsequent rounds of mutagenesis were conducted on an ATCC 17978 *phoU* chromosomal deletion strain, and plated on high phosphate media to reduce the background level of cleavage of the chromogenic substrate. Chromosomal deletions were performed by allelic exchange using a pEX2tetRA vector, which was created by insertion of the *tetRA* tetracycline resistance marker from Tn10 into the pEXG2 plasmid **(42)**. Roughly 1000bp regions upstream and downstream of the genes of interest were amplified for homologous recombination with the ATCC 17978 chromosome. Sucrose was used to counter-select against cells retaining the pEX2tetRA backbone, and deletions were confirmed by PCR. Complementation of deletions was accomplished by repairing the original deletion in the chromosome, again using the pEX system and allelic exchange.

Donor *E. coli* containing the pMarKT transposon vector were suspended in LB broth to an OD600 of 40 and mixed with an equal volume of the recipient *A. baumannii* suspended to OD_600_ of 20. 50 μL aliquots of this mixture were then plated in spots on a dried LB agar plate and incubated for 2 h at 37 °C **(41)**. Each 50 μL spot resulted in about 80,000 colonies of *A. baumannii* containing Mariner transposon insertions. The mutants were plated on LB agar containing 1X M63 salts, 50 μg/mL kanamycin, 30 μg/mL chloramphenicol, and 40 μg/mL XP substrate. Plates were incubated for at least 36 h at 30 °C to allow for the appearance of the blue color from cleavage of the XP substrate. Sequencing of the transposon insertions was adapted from the method described in Chun et. al. **(43)**, including semi-arbitrary two-step PCR amplification of transposon regions followed by sequencing.

### Ethidium bromide uptake assay

Bacteria were grown in 5 mL LB cultures to mid-log OD600 (0.3-0.6), then spun down and normalized in PBS to OD600 0.2. Prior to measurement, CCCP was added at 200 μM to inhibit the activity of efflux pumps. Ethidium bromide was added immediately prior to measurement to final concentration of 1.2 μM in 200 μL total reaction volume. Permeability was assessed using a PerkinElmer EnVision 2104 Multilabel Reader using a 531 nm excitation filter, 590 nm emission filter, and a 560 nm dichroic mirror. Readings were taken every 15 s for 30 min with samples assessed in triplicate in a Greiner bio-one 96-well flat bottom black plate.

### MIC measurements

MICs were determined in 96-well microtiter plates using a standard two-fold broth dilution method of antibiotics in LB broth. The wells were inoculated with 10^4^ bacteria per well, to a final well volume of 100 μL, and plates were incubated at 37 °C with shaking unless stated otherwise. Experiments were performed thrice using two technical replicates per experiment. MICs were interpreted as the lowest antibiotic concentration for which the average OD600 across replicates was less than 50% of the average OD_600_ measurement without antibiotic.

### Crystal violet assay for exopolysaccharide production

Strains were inoculated to OD_600_ 0.05 and grown overnight at 37 °C in 2 mL LB broth with shaking in glass tubes. The next day, liquid was carefully decanted and the tubes left to dry for 2 h at 37 °C. Pellicles were stained with the addition of 0.1% crystal violet, then gently washed three times in dH_2_O. Crystal violet was solubilized in a 80:20 solution of ethanol:acetone and read at 590 nm. P values were determined from a Student’s t-test over three biological replicates per sample.

### Membrane isolation, GPL extraction, and TLC

Cells were collected at specified time points and spun down at 17,000 g for 10 min. Spheroplast formation and sucrose gradient separation of IM and OM was adapted from a method by Osborn et. al (44) by use of a defined 73%-53%-20% sucrose gradient as described in Dalebroux et. al. (45). The purity of membrane separation by this method was confirmed by Western blotting for the *A. baumannii* OM-localized OmpA protein, with 10 μg of total protein loaded into each lane as measured by Bradford protein assay (Fig. S10). GPLs from isolated membranes were extracted using a 0.8:1:2 ratio of water : chloroform : methanol as per the method of Bligh and Dyer (46). Two-dimensional TLC was performed using silica gel 60 plates and immersion in Solvent System A (60:25:4 CHCl_3_:CH_3_OH:H_2_O), followed by Solvent System B (80:12:15:4 CHCl_3_:CH_3_OH:CH_3_COOH:H_2_O) in the orthogonal direction.

### Cryo-EM sample preparation, data acquisition and image processing

Purified Mla complex at ~1 mg/ml was applied to glow-discharged holey grids, blotted for 6 s, and plunged in liquid ethane using a Vitrobot (FEI). Images were acquired on a FEI Tecnai G2 F20 200 kV Cryo-TEM equipped with a Gatan K-2 Summit Direct Electron Detector camera with a pixel size of 1.26 Å/pixel. 500 micrographs were collected using Leginon (47) spanning a defocus range of -1 to -2 μm.

Movie frames were aligned with MotionCorr2 (48) and the defocus parameters were estimated with CTFFIND4 (49). 333 high-quality micrographs were selected by manual inspection, from which ~55,000 particles were picked with DOG in Appion (50). Particle stacks were generated in Appion using a box size of 200 pixels. Several successive rounds of 2D and 3D classification were performed in Relion 2 (51, 52) using an initial model generated by Common Lines in EMAN2 (53) leading to a final stack of ~ 14,000 particles for 3D structure refinement in Relion.

### Structure modeling and docking in the EM density

The structures of MlaB and MlaF were modeled using the threading server Phyre (54) based on the structures of the anti-sigma factor antagonist tm1081 (PDB ID 3F43, 18% sequence identity to MlaB) and the ABC ATPase ABC2 (PDB ID 1OXT, 36% sequence identity to MlaF) respectively. Two copies of each structural model were positioned in their putative location within the EM map using Chimera (55) and their position was optimized using the Fit to EM map option. The abMlaD and abMlaE structures were modeled on ecMlaD and ecMlaE structural models deposited in the Gremelin database (21), using Modeller. For abMlaE, the hexamer was modelled with Rosetta (56) as described previously (4).

### Membrane Isolation and Separation

Cells were resuspended in 20 mL of 0.5 M sucrose, 10 mM Tris pH 7.8, 75 μg freshly prepared lysozyme (Roche 10837059001), and 20 mL of 0.5 mM EDTA, and kept on ice with gentle stirring for 20 min. Samples were homogenized (Avestin EmulsiFlex-C3) and spun down at 17,000 g for 10 min to removed un-lysed cells prior to ultracentrifugation. Membranes were spun down using a Ti45 Beckman rotor at 100,000 g for 1 h and then added to the top of a sucrose gradient. IM and OM were separated by 18-hour ultracentrifugation using a SW-41 rotor in a Beckman Coulter Optima L90X ultracentrifuge.

### MlaC and MlaD protein purification and GPL extraction

Primers were designed to amplify the *mlaC* gene of ATCC 17978, excluding the signal sequence for export from the cytoplasm, and the periplasmic domain of *mlaD* of ATCC 17978, excluding the membrane-spanning domain. These fragments were cloned into pET29b and expressed with a carboxy-terminal hexahistidine (-6HIS) tag in BL21 *E. coli* with 2 h induction. Cells were pelleted and resuspended in Tris-buffered saline containing 10% glycerol (TBSG) and protease inhibitor cocktail (Roche, Complete EDTA-free). Cells were lysed by homogenization (Avestin) and ultracentrifuged at 100,000 g for 1 h to spin down membranes. The supernatants were then applied to a 5 mL-HiTrap(TM) Chelating HP Ni-affinity column pre-loaded with 0.1 M NiSO_4_ and equilibrated with TBSG. The proteins were eluted from the column using FPLC (Akta) by applying a stepwise gradient of 25 mM, 50 mM, and finally 300 mM imidazole for protein elution. Elution was monitored by UV-absorption at 280 nm. The MlaC- and MlaD-containing fractions were then further purified by injecting into a HiLoad 120 ml-6/600 Superdex(TM) 200 preparative grade size-exclusion column equilibrated in TBSG using a flow rate of 1 mL/min. The purity of the collected protein fractions was confirmed by SDS polyacrylamide gel electrophoresis. Proteins were diluted to 2 mg/mL and dialyzed overnight in 1 L TBSG at 4 °C with stirring. GPLs were extracted from 1 mg each of purified proteins MlaC and MlaD by the method of Bligh and Dyer and analyzed by LC-MS/MS as previously described.

### LC-MS/MS

Retention of PG, CL, PE, and Lyso-CL was achieved at a flow rate of 0.3 mL/min using mobile phase A [CHCl_3_/CH_3_OH/NH_4_OH (800:195:5 v/v/v)] and mobile phase B [CHCl_3_/CH_3_OH/NH_4_OH (600:340:5 v/v/v)]. The chromatography method used is a three-step gradient as described in the SI Materials and Methods of Dalebroux et al. (26). The samples were run on an Agilent Zorbax Rx-SIL silica column (2.1x100mm, 1.8-Micron) using an Agilent HPLC autosampler. Mass spectrometry was performed using an AB Sciex API4000 Qtrap with multiple reaction monitoring (MRM). The identities of the major GPLs present in the *A. baumannii* membrane were predicted by parent ion scans.

### Stable isotope assay development

The Q1/Q3 transitions of glycerolphospholipids from cells grown in 2-^13^C acetate were determined using a Thermo Orbitrap LTQ. The integrated peak areas of both ^13^C-labeled and unlabeled GPLs from the AB Sciex API4000 Qtrap were used to calculate the ion-current ratios for each GPL species. The ratio of labeled GPL for each unique species can be calculated based on the following equation:

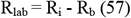

Where R_i_ is the ion-current ratio of labeled GPL to unlabeled GPL within the sample and R_b_ is the ion-current ratio of samples before the administration of the tracer, ^13^C-acetate, and represents the natural background abundance of the stable isotope species within the bacterial membrane. R_lab_ approximates the molar ratio of labeled species to unlabeled species (n_lab_/n_un_) according to the equation (n_lab_/n_un_) = [R_i_-R_b_]/k, where k is the molar response factor of the instrument and is ideally equal to unity (57).

To demonstrate that OM phospholipases will not distinguish between labeled and unlabeled GPL and therefore will not affect the ratio of labeled to unlabeled GPL obtained from this assay, we compared ratios of labeled and unlabeled GPL from wild type *A. baumannii* and deletion mutants in *pldA*. Bacteria were grown carrying either the empty pMMB::*kan* vector, or expressing the Walker box mutant MlaF-K55L. Accumulation of newly synthesized GPL was observed in those strains expressing MlaF-K55L when compared to the vector control, across various species of GPL. Of strains expressing the vector control, on average 51.84% ± 1.07% and 52.07% ± 1.23% of newly synthesized PG C16:0/18:1 appeared on the inner membrane of wild type and Δ*pldA*, respectively, after one hour incubation with ^13^C acetate, while 66.33% ± 1.23% and 62.60% ± 1.70% of newly synthesized PG C16:0/18:1 accumulated at the inner membranes of wild type and Δ*pldA* expressing MlaF-K55L. In vector controls strains, 48.53% ± 1.37% and 51.01% ± 0.55% of newly synthesized PG C16:0/16:0 appeared on the inner membrane of wild type and Δ*pldA*, respectively, after one hour incubation with ^13^C acetate, while 62.98% ± 1.01% and 60.41% ± 1.25% of newly synthesized PG C16:0/16:0 accumulated at the inner membranes of wild type and Δ*pldA* expressing MlaF-K55L. In vector controls strains, 50.17% ± 1.31% and 50.49% ± 1.15% of newly synthesized PE C16:0/18:1 appeared on the inner membrane of wild type and Δ*pldA*, respectively, while 60.14% ± 0.93% and 62.06% ± 1.07% of newly synthesized PE C16:0/18:1 accumulated at the inner membranes of wild type and Δ*pldA* expressing MlaF-K55L.

### Stable isotope GPL analysis and culture conditions

Cultures of *A. baumannii* ATCC 17978 were grown in M63 media containing 5 mM sodium acetate and 4 mM MgCl_2_ to OD600 0.4, then washed and resuspended in media containing 5 mM 2-^13^C sodium acetate (Cat. No. CLM-381-0, Cambridge Isotope Laboratories, Inc.). Membrane fractions were isolated from both wild type and *mla* mutant *A. baumannii* at simultaneous time points, and GPL were extracted and assessed using previously established LC-MS/MS methods with additional MRM values to account for the increased m/z ratios of ^13^C-labeled GPL. MRMs were selected to account for PG and PE having acyl chains of either C16:0/16:0, C16:0/18:1, and C18:1/18:1 as these were determined by total ion scan MS to be the predominant species of PG and PE GPL. Pulse experiments were performed at least twice for each mutant. Further details of assay development are described in SI Materials and Methods.

## Acknowledgements

We thank Dale Whittington and Dr. Scott Edgar at the Mass Spectrometry Center, Department of Medicinal Chemistry, University of Washington for technical help with MS analysis; and Mauna Edrozo for technical help.

## Author Contributions

CK carried out all microbiology experiments, purified membranes and lipids and analyzed them by mass spectrometry, and wrote the paper with SIM. JF purified the Mla protein complex, analyzed its components and wrote that section of the paper. HDK participated in bioinformatic analysis of Mla and experimental design. ZDD advised as to the mass spectrometry lipid analysis and participated in data analysis. JB performed electron microscopy analysis of the protein complex and wrote that section of the paper. AB and JMK performed electron microscopy analysis of the protein complex. SIM planned and supervised all the experiments, developed the genetic screen for OM permeability, analyzed data, and wrote the paper with CK.

## Author Information

Authors have no competing financial interests. Correspondence and Requests for materials should be addressed to SIM.

## Data Availability

The cryo-EM map has been deposited in the Electron Microscopy Data Bank with accession code EMD-8738 (8.7 Å map). The coordinates for the MlaBDEF model have been deposited to the PDB-dev database.

---

**Fig. S1.**
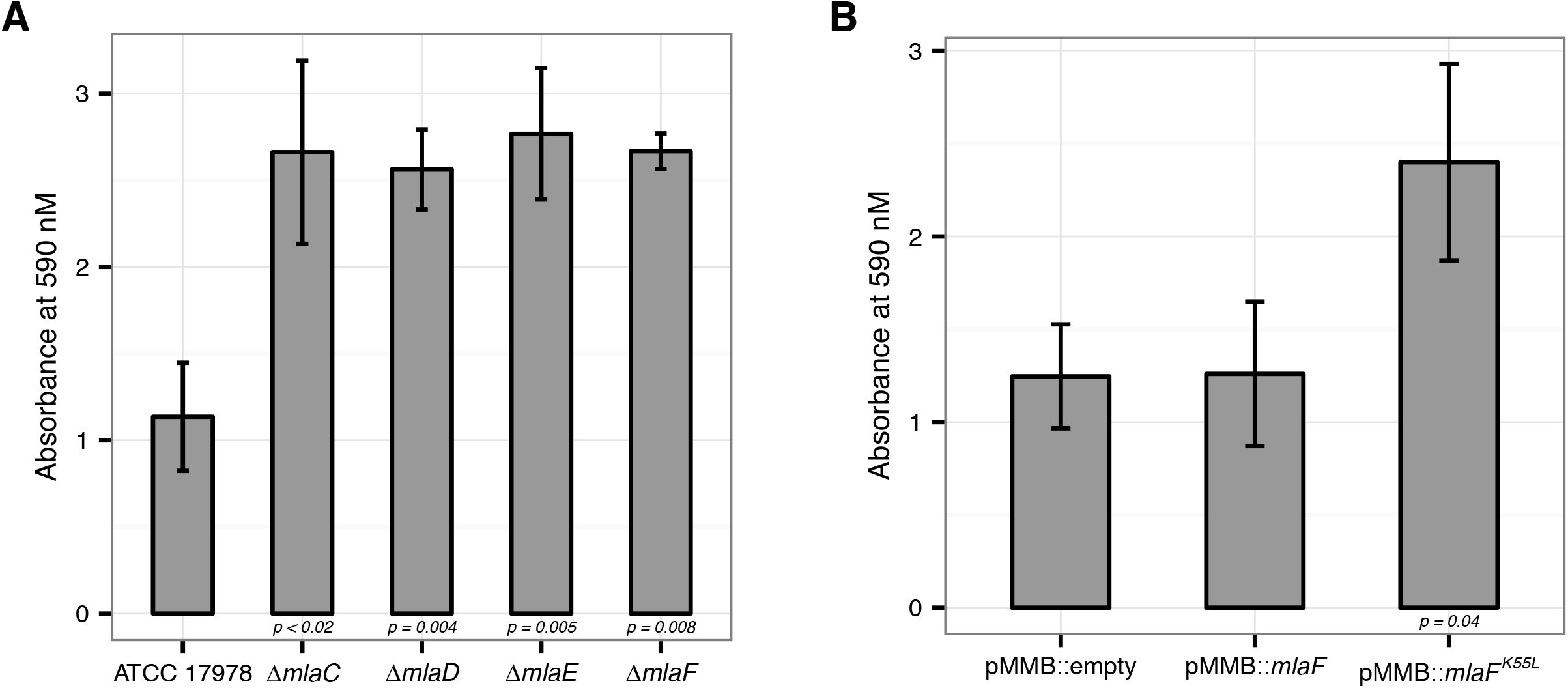
Disruption of the Mla system leads to an increase in exopolysaccharide production. **(A)** Quantification of crystal violet staining from *mla* deletion mutants. Error bars represent ± s.d. for biological replicates (n = 3). **(B)** Quantification of crystal violet staining following plasmid expression of MlaF compared to its dominant negative version, MlaF^K55L^. Error bars represent ± s.d. for biological replicates (n = 3).

**Fig. S2.**
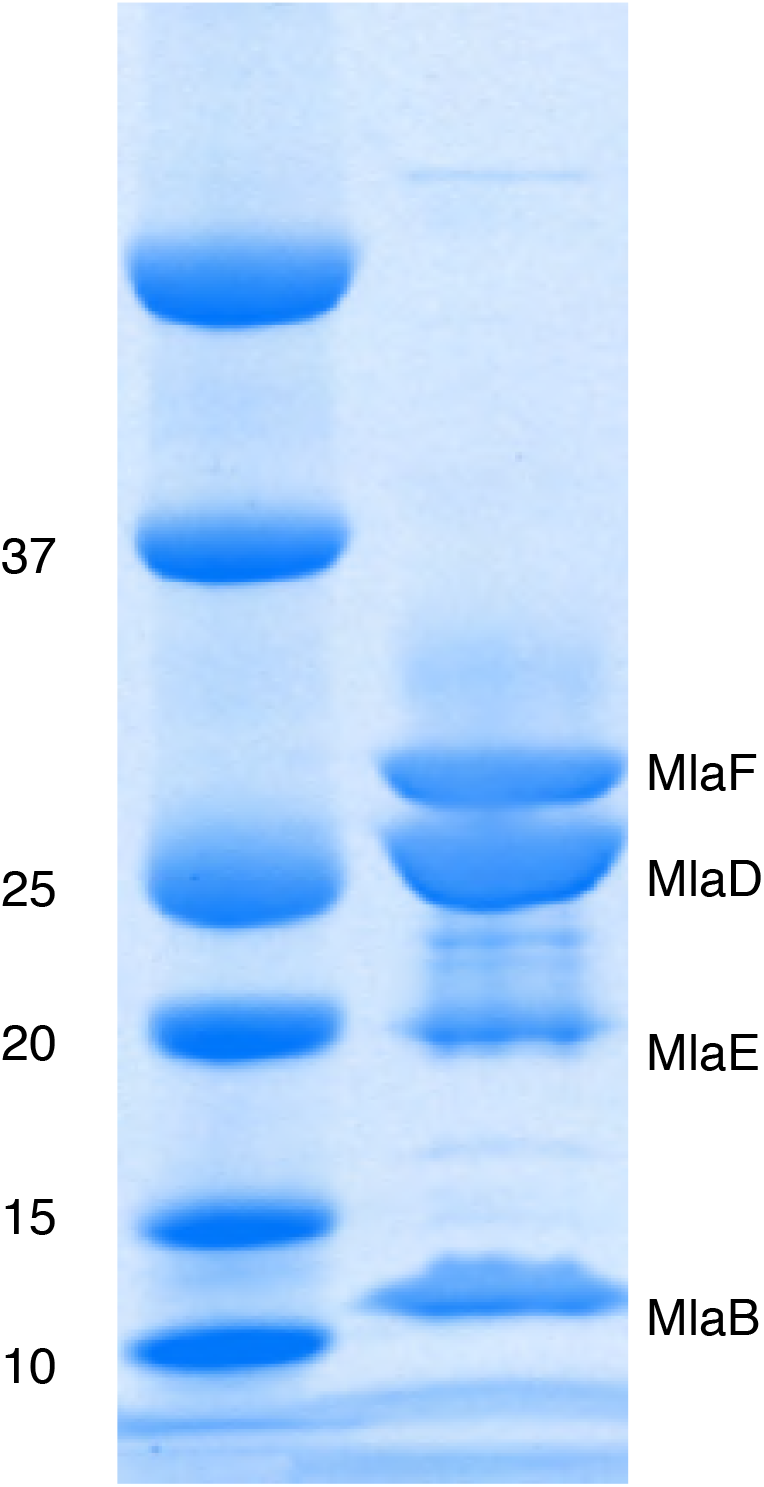
Mla components copurify following protien expression. SDS-PAGE analysis of proteins copurified with hexahistidine-tagged MlaB (-His6). Band identities were assigned based on MS.

**Fig. S3.**
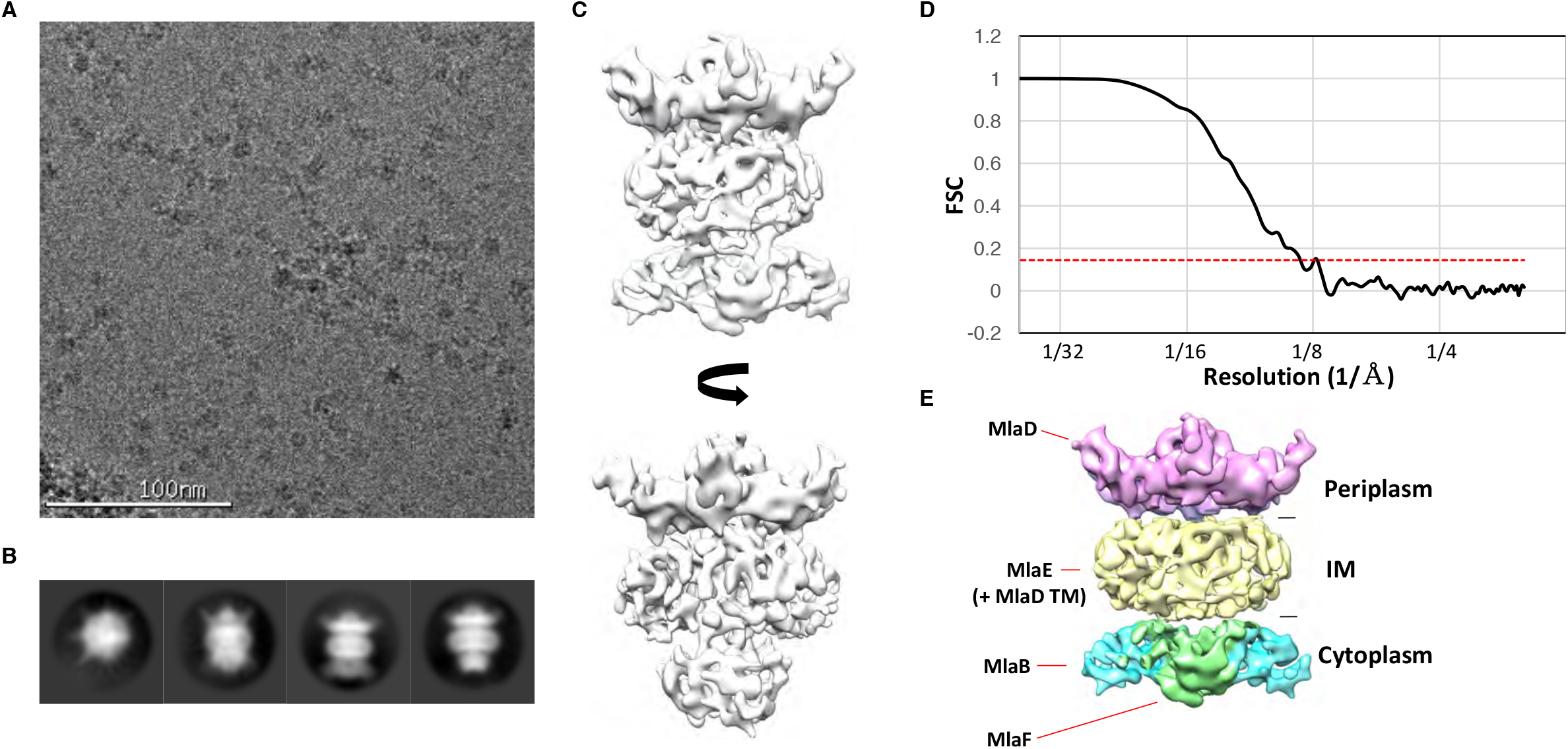
Cryo-EM structure of the abMlaBDEF complex. **(A)** Representative electron micrograph region of frozen-hydrated MlaBDEF complex. The scale bar is in white atthe bottom. **(B)** Representative reference-free 2D class averages of abMlaBDEF, generated using Relion, illustrating the various views observed. **(C)** Cryo-EM map of the Mla complex, shown in two orientations corresponding to the two last classes shown in B. **(D)** The FSC curve for the MlaBDEF structure is shown in black, with the gold-standard resolution definition of 0.143% indicated with a red dotted line. The nominal resolution for this stracture is 8.7 Å. **(E)** The regions of density attributed to the periplasmic domain of MlaD, the TM domains of MlaD and MlaE, and to MlaB and MlaF are in pink, yellow, cyan and green respectively.

**Fig. S4.**
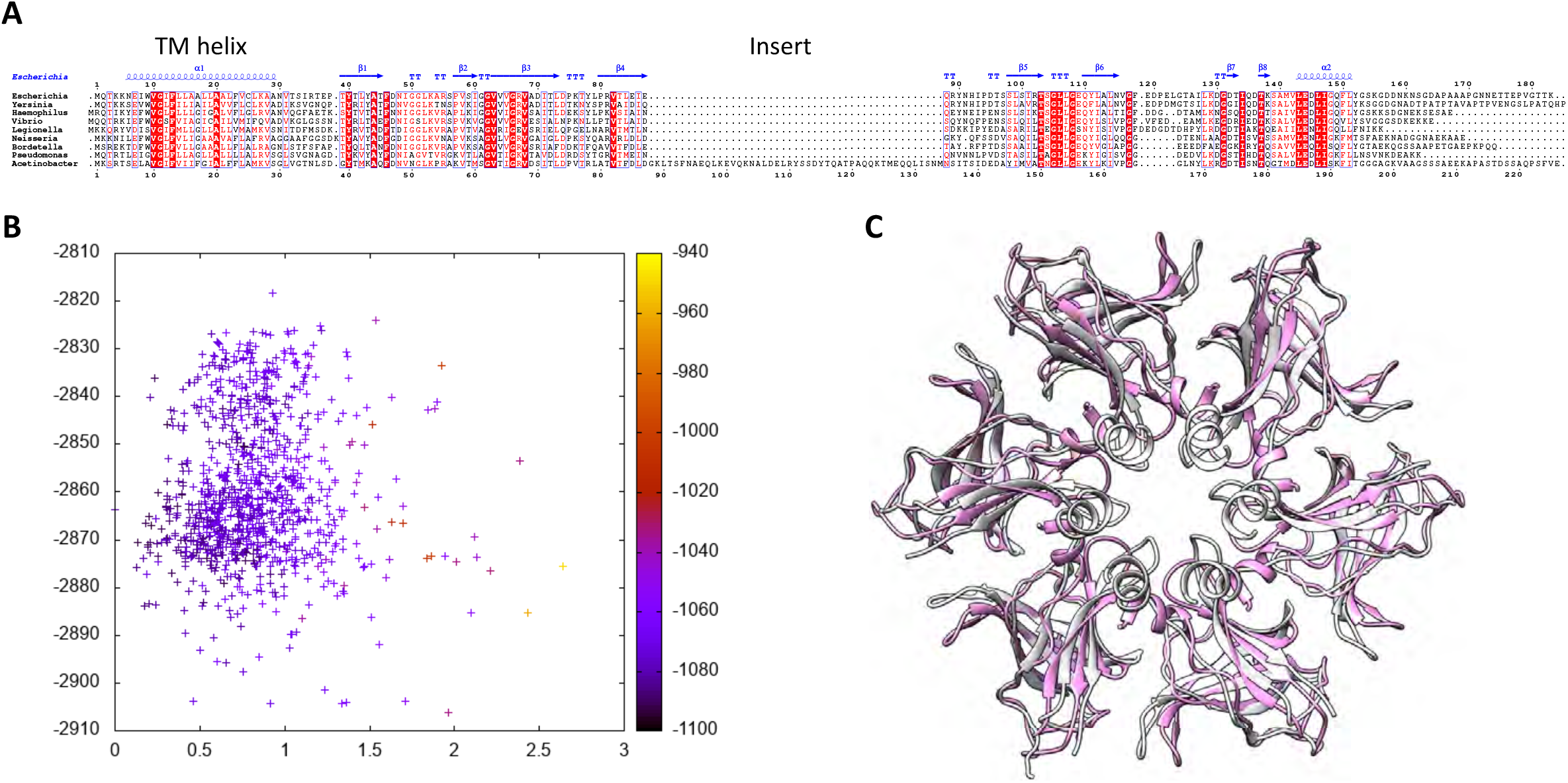
Modeling of the abMlaD hexamer. **(A)** Multiple alignment of MlaD sequences from various gram-negative human pathogens. The secondary structure for ecMlaD is shown at the top. The position of the abMlaD insert is indicated. **(B)** Result of the all-atom refinement step for the MlaD hexameric model. The energy of each model is plotted versus the RMSD relative to the initial model, and color-coded for the fit to the EM map density. **(C)** Cartoon representation of our abMlaD hexameric model (magenta), superimposed to the crystallographic ecMlaD hexamer structure (grey).

**Fig. S5.**
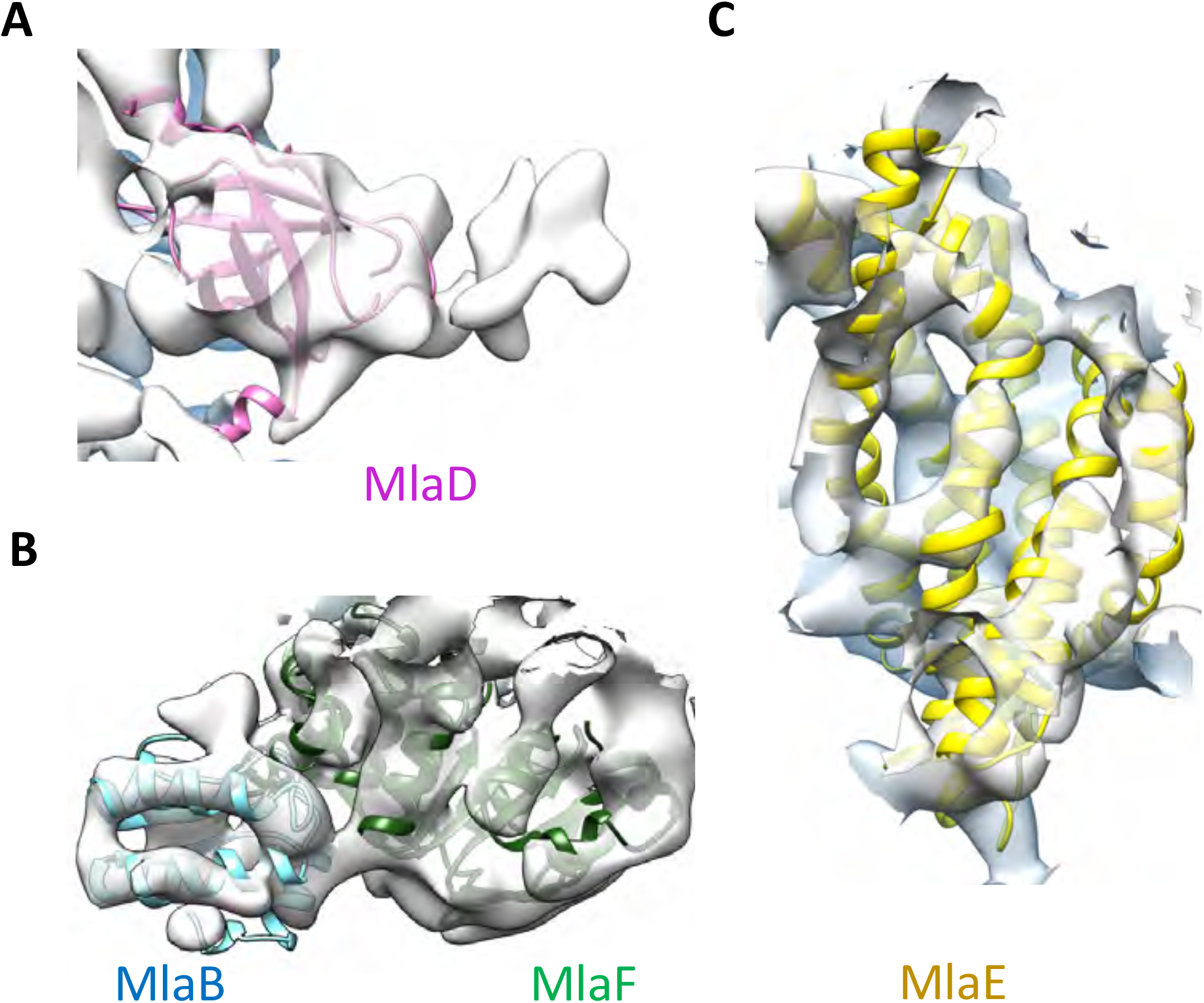
Close-up view of the MlaB, MlaD, MlaE and MlaF models in the abMlaBDEF cryo-EM map. **(A)** Region of the density corresponding to a MlaD monomer, with the corresponding atomic model in magenta. **(B)** Region of the density corresponding to a MlaF-MlaB hetero-dimer, with the corresponding atomic models in green and cyan respectively, shown from two different angles. Density for helices are well defined for most of the model. **(C)** Region of the density corresponding to a MlaE monomer, with the corresponding atomic model in yellow.

**Fig. S6.**
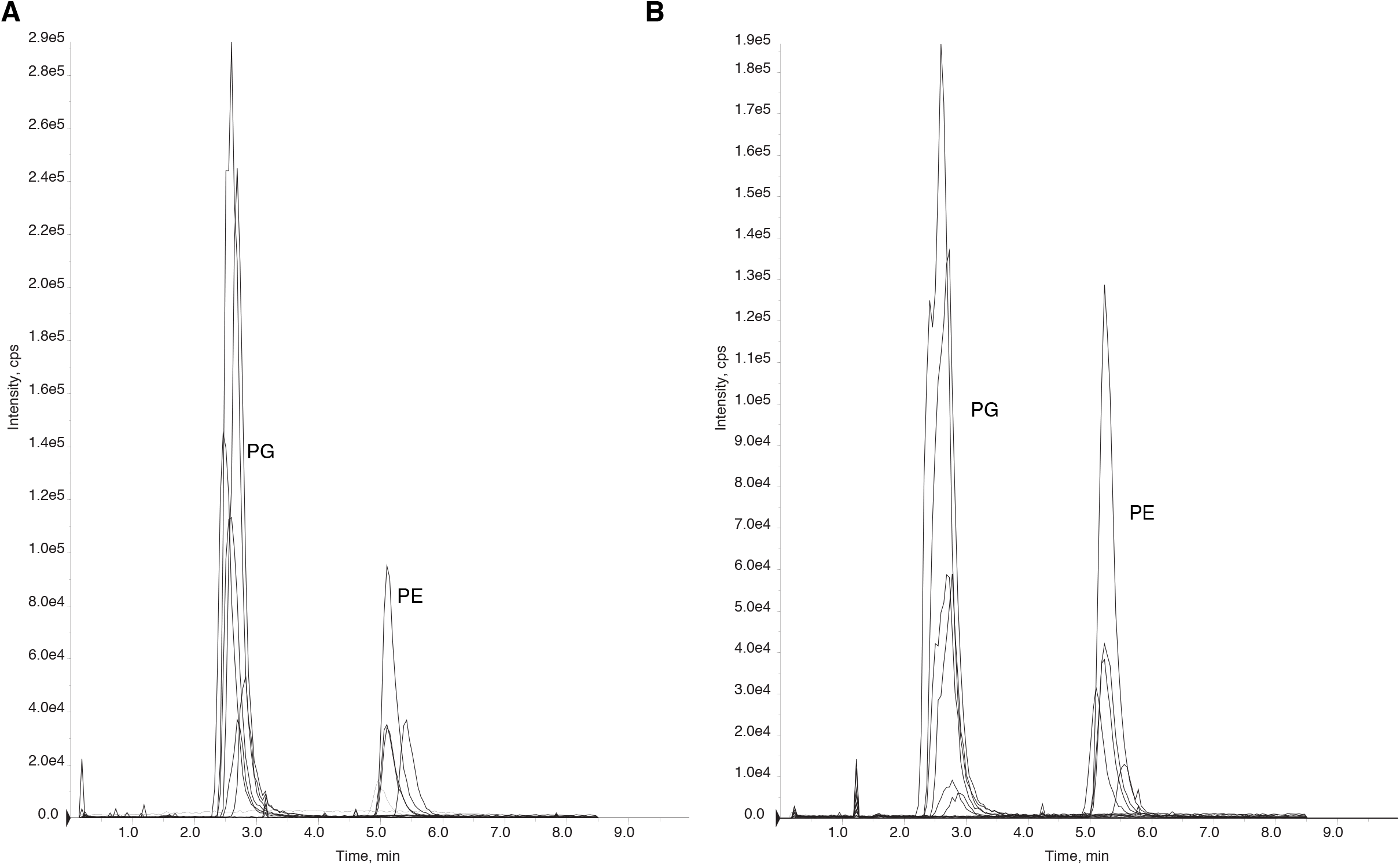
Purified periplasmic components of the Mla system remain bound to glycerophospholipids. **(A)** Chromatogram of LC-MS/MS of glycerophospholipids extracted from purified MlaC. Peaks were identified based on MS and elution time. PG, phosphatidylglycerol. PE, phosphatidylethanolamine. **(B)** Chromatogram of LC-MS/MS of glycerophospholipids extracted from purified MlaD soluble domain.

**Fig. S7.**
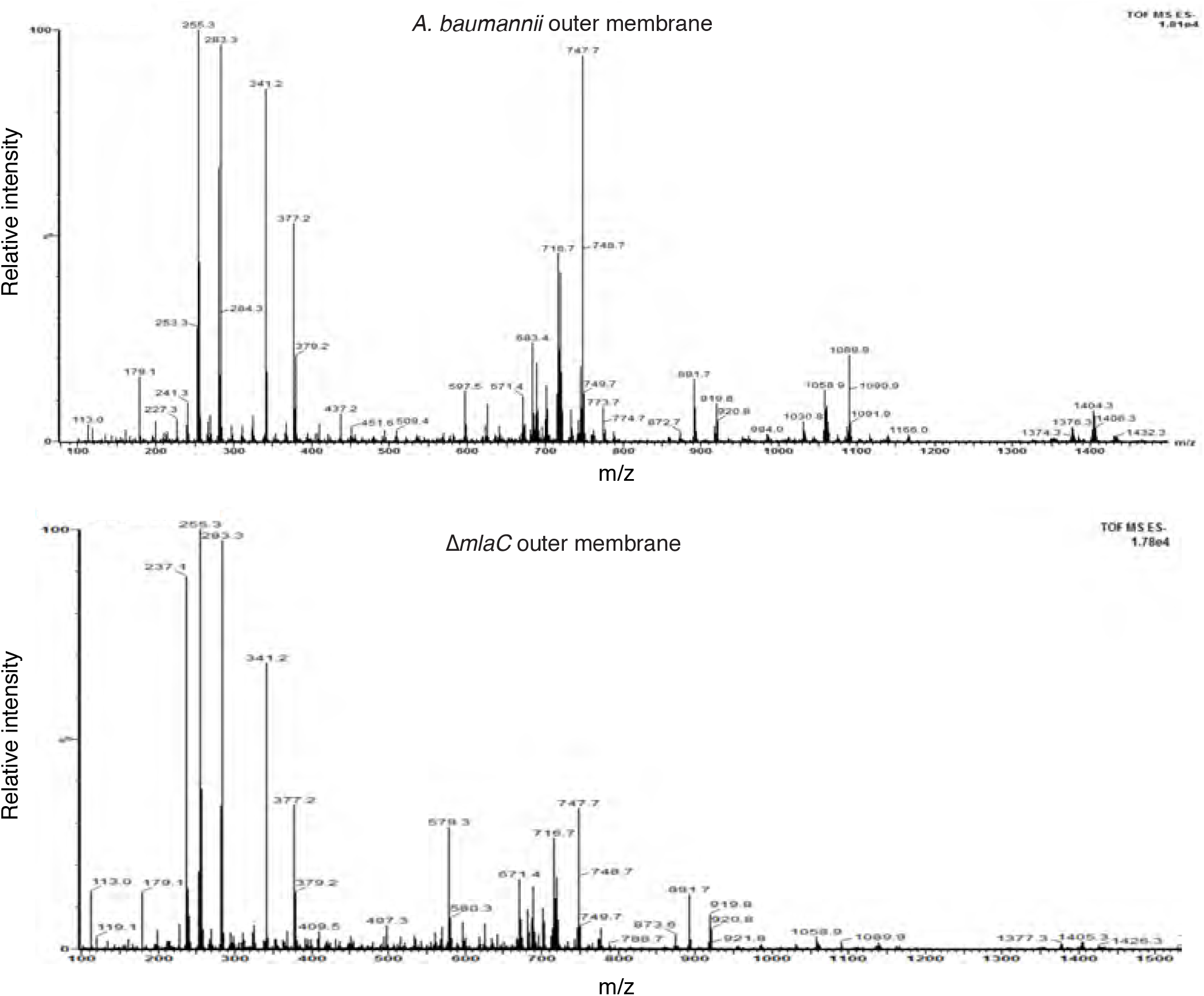
Deletion of *mlaC* results in a reduction in levels of outer membrane glycerophospholipids. Total ion scan of isolated outer membranes of *A. baumannii* and Δ*mlaC*. Typical membrane glycerophospholipids fall within the m/z range of 600-1500.

**Fig. S8.**
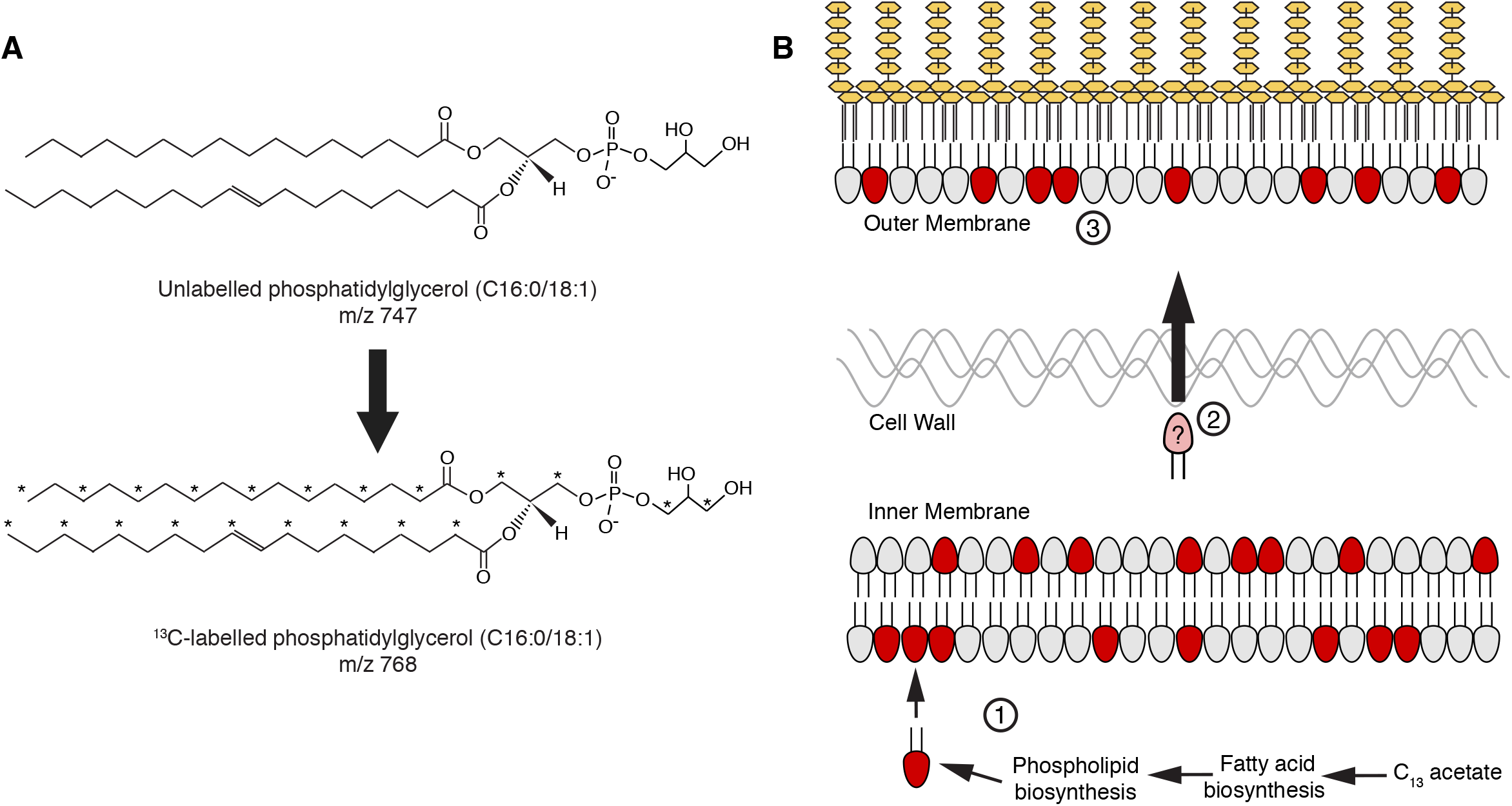
A stable isotope assay of glycerophospholipid transport from the inner membrane to the outer membrane. **(A)** Schematic showing an example of the shift in mass-to-charge ratio (m/z) of glycerophospholipids (GPL) following growth in 2-^l3^C acetate. **(B)** A schematic illustrating the rationale of the stable isotope assay: (1) Newly synthesized ^13^C-labeled GPL, shown here in red, are first inserted into the inner membrane (IM) following synthesis; (2) the likelihood that a given GPL that is trafficked from the IM to the OM will be labeled is proportional to the ratio of labeled to unlabeled GPL in the IM; (3) a comparison of the ratios of labeled to unlabeled GPL in the inner and outer membranes will therefore reflect the efficiency of GPL transport.

**Fig. S9.**
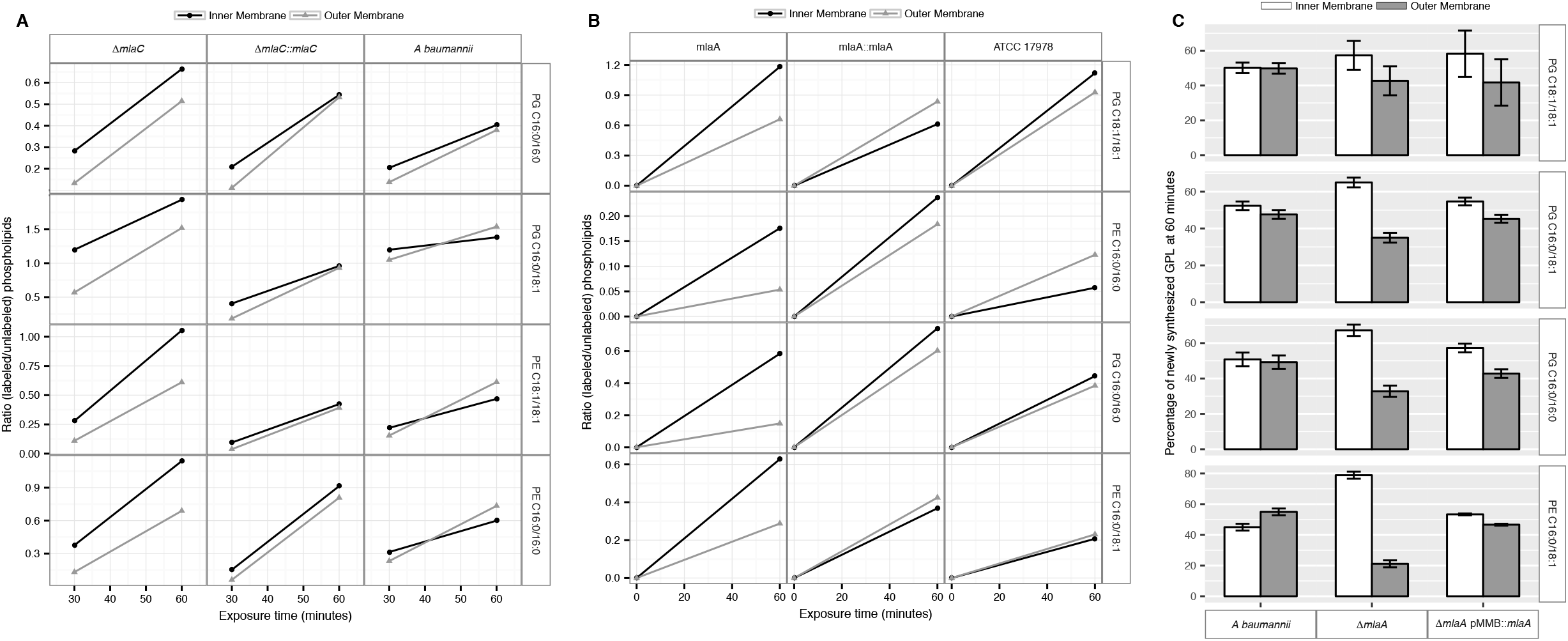
Newly synthesized glycerophospholipids accumulate at the inner membrane of MlaA mutants. **(A)** LC-MS/MS quantification of ^l3^C labelled/unlabeled glycerophospholipids in isolated membrane fractions over time after growth in 2-^l3^C acetate in Δ*mlaC* and complemented strain. Facet labels on the right indicate the specific glycerophospholipid species analyzed and the acyl chain length. PG, phosphatidylglycerol; PE, phosphatidylethanolamine. Shown is representative data from repeated experiments. **(B)** LC-MS/MS quantification of ^13^C labelled/unlabeled glycerophospholipids in isolated membrane fractions over time after growth in 2-^13^C acetate in ΔmlaF and complemented strain. Facet labels on the right indicate the specific glycerophospholipid species analyzed and the acyl chain length. PG, phosphatidylglycerol; PE, phosphatidylethanolamine. Shown is representative data from repeated experiments. **(C)** Relative proportion of newly synthesized GPL on IM and OM after one hour growth in 2-^13^C acetate. Error bars represent ± s.d. (n = 2). Statistical analyses performed using a Student’s t test. p-Value: *, p < 0.05; **, p < 0.01.

**Fig. S10.**
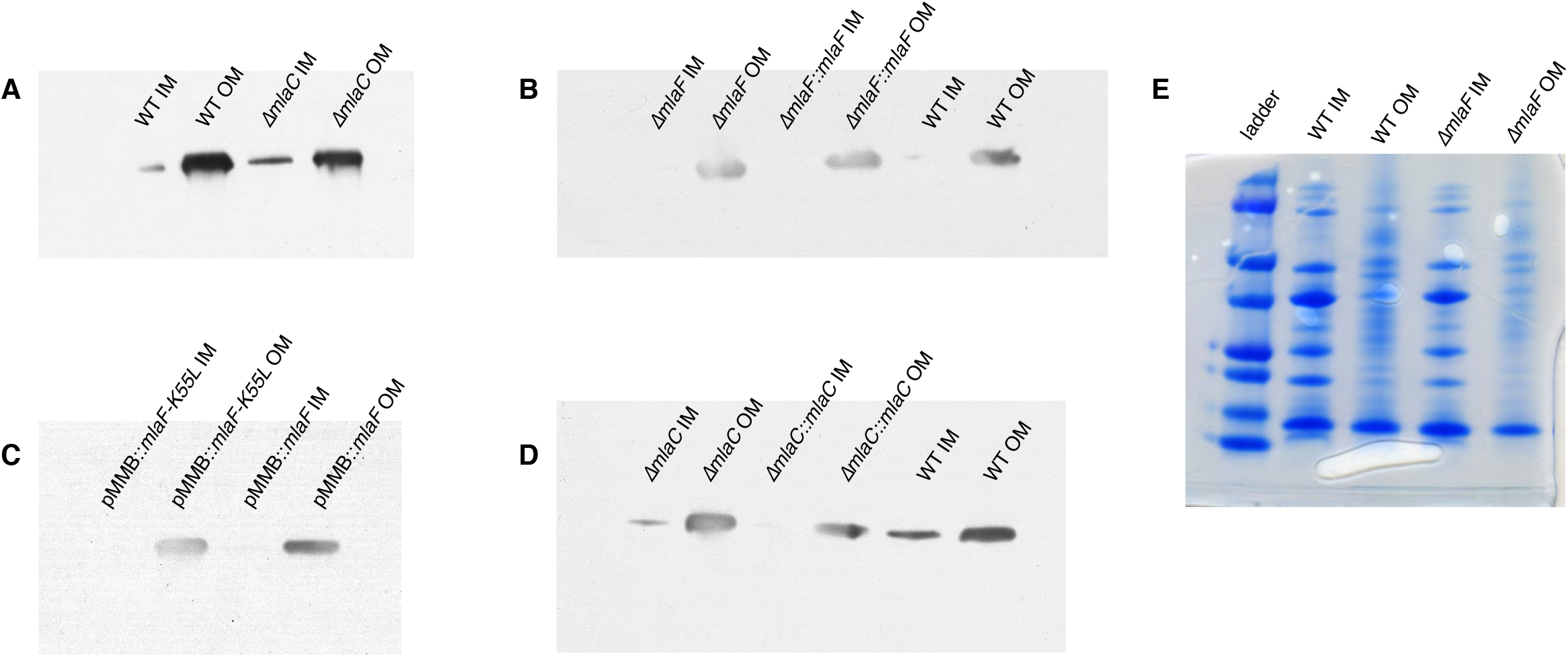
Confirmation of inner and outer membrane separation. Each lane contains 10 μg total protein as measured by Bradford protein assay. **(A)** α-OmpA Western blot of separated membranes analyzed in Fig. 7. **(B)** α-OmpA Western blot of separated membranes analyzed in Fig. 9A **(C)** α-OmpA Western blot of separated membranes analyzed in Fig. 9B. **(D)** α-OmpA Western blot of separated membranes analyzed in Fig. 9C. **(E)** Coomasie stained SDS-protein gel of representative isolated membrane samples alongside BioRad Precision Plus Protein Standard.

**Fig. S11.**
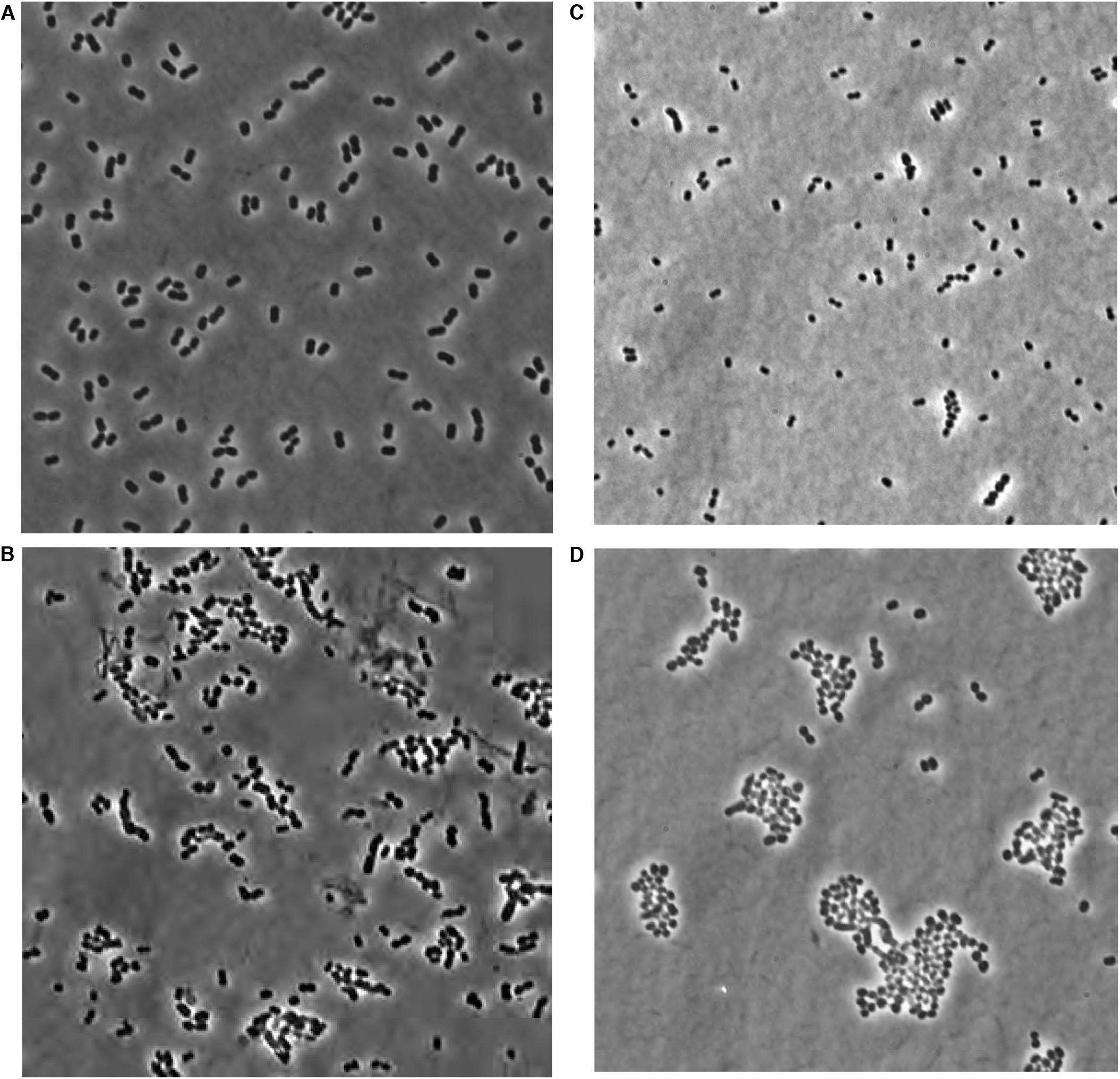
Phase microscopy images of wild type and *mla* mutant *A. baumannii*. Images were collected from cultures grown to mid-log (OD600 0.4-0.6) growth phase from wild type **(A)**, *mlaA* **(B)**, *mlaC* **(C)**, and *mlaF* **(D)** deletion mutants of *A. baumannii*.

**Table S1:**
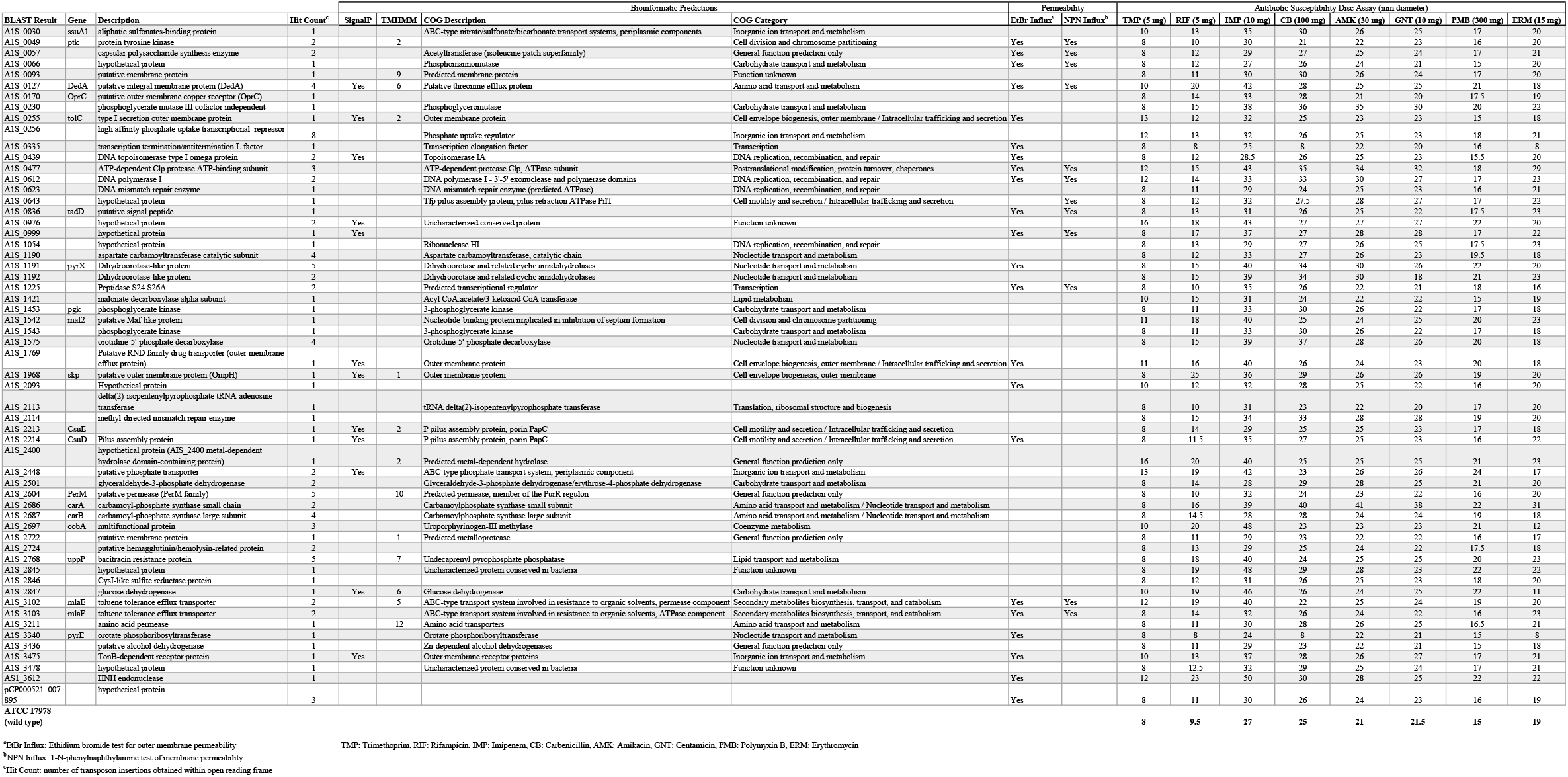
Results of transposon mutagenesis screen for genes involved in outer membrane barrier function in ATCC 17978.

**Table S2:**
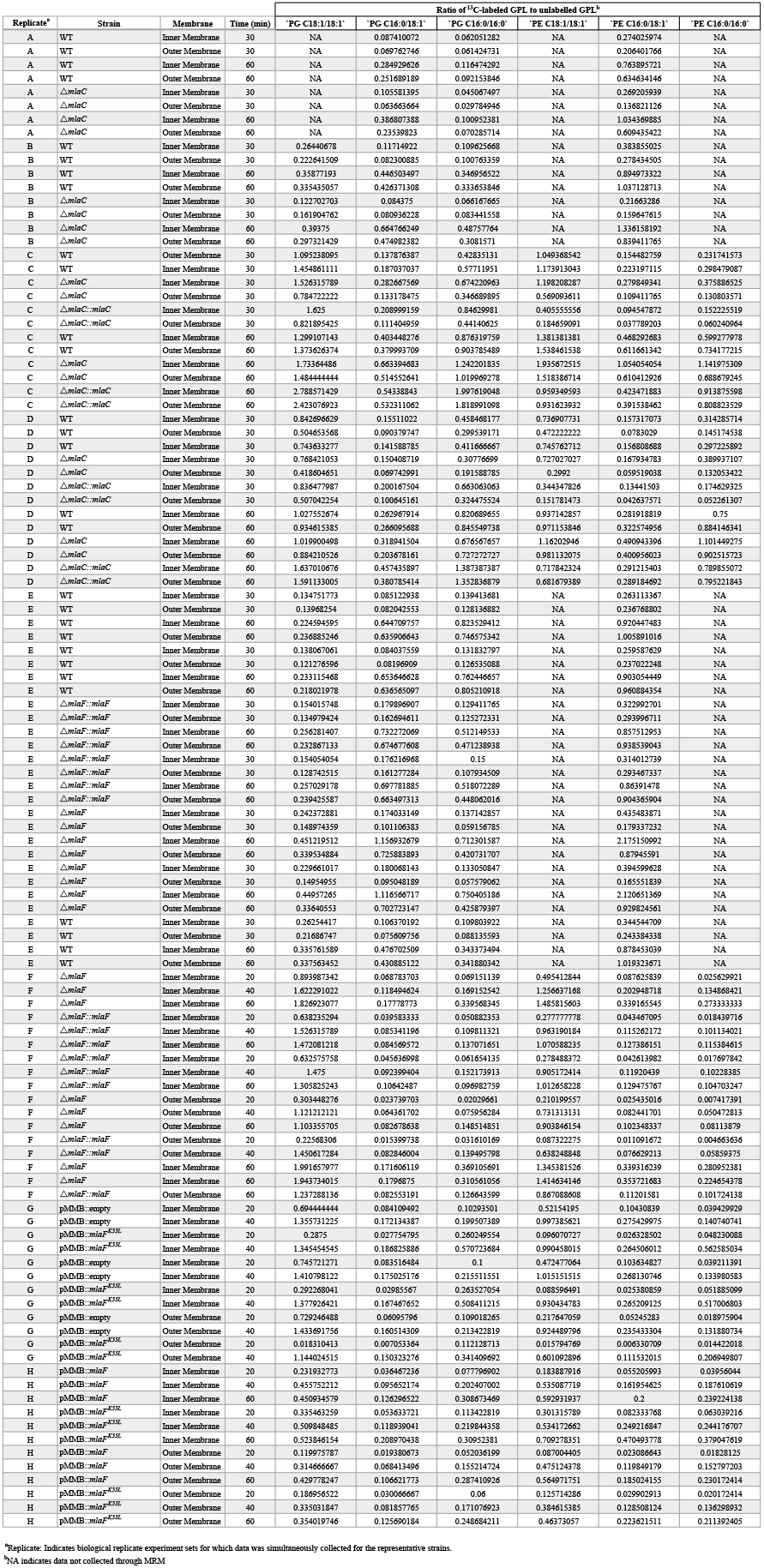
Accumulation of newly synthesized glycerophospholipids in *A. baumannii* inner and outer membranes.

